# Characterisation of guided entry of tail-anchored proteins in *Magnaporthe oryzae*

**DOI:** 10.1101/2025.02.28.640736

**Authors:** Felix Abah, Qiaojia Zheng, Xinru Chen, Linwan Huang, Xiaomin Chen, Jules Biregeya, Osakina Aron, Zonghua Wang, Wei Tang

## Abstract

Rice (*Oryza sativa L.*) is one of the most important staple foods for human population worldwide. However, rice production continues to be severely threatened by rice blast disease caused by an ascomycete fungus *Magnaporthe oryzae*. Tail-anchored (TA) proteins are conserved across diverse organisms, and belongs to a class of polypeptides that are inserted into the membrane by a hydrophobic sequence located at the C-terminal region. The Guided Entry of Tail-anchored (GET) complex is responsible for the post-translational insertion of nascent TA proteins into the ER lipid bilayer. In *Saccharomyces cerevisiae,* the GET pathway comprises of six known associated components Get1, Get2, Get3, Get4, Get5, Sgt2 and Ssa1 that have been identified and extensively studied. However, the role of the GET complex in rice blast fungus has not been elucidated. Here in, we identified five proteins of the GET Complex in *M. oryzae* namely MoGet1, MoGet2, MoGet3, MoGet4 and MoSgt2 and generated the gene knock-out mutants. Deletion of MoGET1 and MoGET2 revealed that they are required for vegetative growth, asexual reproduction and pathogenesis, while MoGet3 negatively regulates hyphal growth, asexual development and pathogenesis of *M. oryzae*. In contrast, loss of MoGet4 and MoSgt2 had no effect on the normal development of the rice blast fungus. We demonstrated that the MoGet2 is important in osmotic stress response while MoGet2 and MoGet3 positively and negatively regulate cell wall integrity, respectively. The MoGet1 and MoGet2 were ER-localised and indispensable for DTT-induced ER stress response. *In vitro* and *in vivo* interaction assay revealed MoGet3 has physical interaction with both MoGet1 and MoGet2 indicating the existence of possible synergistic function amongst the Get components in rice blast fungus. In summary, this finding indicates the GET Components offers an excellent target for the development of antifungal components against recalcitrant plant fungal pathogens.

## Introduction

Rice (*Oryza sativa L.*) production generates income and employment for more than 200 million households around the world [1], especially for low-income earners in rural and semi-urban areas in Africa, Asia, Latin America and the Caridean [2]. Total rice consumption is expected to increase to around 584 million tonnes by 2050 [3]. However, rice production is severely affected by rice blast disease, caused by the filamentous fungus *Magnaporthe oryzae* (anamorph: *Pyricularia oryzae*). Rice blast is a widespread disease that affects rice production around the world and therefore poses a threat to global food security [4]. The annual yield loss due to rice blast are around 30% worldwide [4–6], which corresponds to a quantity that can feed 60 million people [7, 8]. In addition, *M. oryzae* causes blast disease in other grass species, including barley and wheat (*Triticum aestivum*) [8, 9].

To initiate blast disease, the three-celled, teardrop-shaped conidium of *M. oryzae* attaches firmly to the surface of the host (with the help of strong, glycoprotein-rich mucilages) [10] and germinates under favorable conditions [11]. A highly melanized and turgor-pressurized dome-shaped infection cell, the appressorium, then forms at the tip of the germ tube. A penetration peg emerges from the dome-shaped appressorium, which penetrates the plant cuticle and cell wall and branches into invasive hyphae [4, 11]. A mature appressorium is highly melanized and chitin-rich, and accumulates glycerol to generate about 8.0 MPa turgor pressure used to mechanically breach the hydrophobic host surface [12].

Tail-anchored (TA) proteins are a class of proteins possessing C-terminal hydrophobic trans-membrane domains [13], and are conserved across diverse organisms, including bacteria, *S. cerevisiae*, *Homo sapiens, Toxoplasma gondii* and plants [14]. They share a topology of cytosolic N-terminal region and a transmembrane domain (TMD) at their C termini [14–16]. However, they lack N-terminal signal peptide, and are therefore targeted to the membrane by posttranslational mechanisms [17]. They are implicated in determining organellar identity and mostly play fundamental roles in cellular metabolism and organismal survival [17–19]. However, malfunction of these proteins may result in disease and aging [20].

Generally, unlike soluble proteins, newly synthesized integral membrane proteins face several challenges, such as aggregation or interaction with other proteins inappropriately in the cytosol, misfolding resulting in non-functional or potentially harmful protein structures and mislocalization to non-intended destinations leading to protein dysfunction [21]. These phenomena occur as the nascent proteins traverse the aqueous cytosolic environment before reaching their membrane destination [21] and pose acute challenges to protein homeostasis in the living cell [22].

To address these challenges, cells have evolved various mechanisms to ensure the proper biogenesis and trafficking of integral membrane proteins [14]. These mechanisms involve the assistance of molecular chaperones, signal sequences, and protein translocation machinery. This compartmentalization and protein quality control help minimize the risk of misfolding, aggregation, and mislocalization during the biogenesis of integral membrane proteins.

In the past decades, different protein-delivering pathways that deliver proteins to different organelles such as ER, mitochondria, chloroplast and peroxisomes have been extensively studied and described. The sorting of TA proteins through these different pathways is guided by physicochemical properties such as length and hydrophobicity index of TMDs, and charge of C-terminus, but not by motif sequence [14, 17].

The Guided Entry of the Tail-Anchored Protein (GET) Pathway is highly conserved in all eukaryotic organisms [19, 23] and functions to post-translationally integrate TA proteins into the membrane of the ER [24, 25]. The GET pathway has about 6 (six) known associated component factors for the recognition, shielding, trafficking and insertion of newly synthesized protein substrate into the ER lipid bilayer. In budding yeast, transmembrane domain (TMD)-recognition complex (TRC) Get4 recruits Get3, Get5 recruits Sgt2, and Sgt2 can recruit other chaperones [19, 25–27]. Get3 transfer TAs to Get1/Get2 insertase in the ER lipid bilayer. The pathway was first identified and studied in yeast, followed by mammals where work on a model animal, *Mus musculus*, identified a 40-KDa ATPase, Transmembrane Recognition Complex (TRC40) [21]. TRC40 is the homolog of Get3 (guided entry of tail-anchored proteins factor 3, ATPase) in yeast [28].

In rice blast fungus, neither the Ssa1 protein nor the GET/TRC pathway homologs have been theoretically identified or experimentally validated. Therefore, this study first identified the *M. oryzae* homolog of Get1, Get2, Get3, Get4 and Sgt2 based on the sequence, structure and functional similarity to the Gets of *S. cerevisiae* origins. However, we could not identify the *M. oryzae* homolog of the Get component, Get5/UBL4A. Nonetheless, we believe this is the first study that provides the functional characteristics of GET machinery in rice blast fungus, *M. oryzae,* and will offer new potential for disease targeting/biotechnological rice breeding programme and future investigation into the dynamic of this pathway and its development for disease intervention.

## Materials and methods

### Fungal strain, plant and culture condition

The WT Guy11 (obtained from the Fungal Genetics Stock Centre, FGSC 9462) stain was used for the generation of the deletion mutants namely (Δ*Moget1*, Δ*Moget2*, Δ*Moget3*, Δ*Moget4* and Δ*Mosgt2*). Plants for infection assays included the susceptible rice cultivar (*O. sativa* CO-39) and barley cultivar, Golden Promise.

The WT Guy11, mutants and the complemented strains were cultured in the complete medium (CM: 6 g yeast extract, 6 g casein hydrolysate and 10 g sucrose in 1 L ddH2O) and complete medium II (CM II: 50 mL 20× Nitrate salt, 1 mL 1000× trace elements, 1 mL1000× vitamin solution, 10 g D-glucose, 2 g peptone, 1 g casein hydrolysate, 1 g yeast extract and 15 g agar powder in 1 L ddH_2_O). Other media used include oatmeal agar (OA: 40 g of oatmeal granules and 20 g of agar powder in 1 L of ddH_2_O), rice straw decoction (SDC: 100 g rice straw and 20 g agar powder in 1 L of ddH_2_O), rice bran (RB: 40 g rice bran and 20 g agar powder in 1 L of ddH_2_O), minimal medium (MM: 6 g NaNO3, 0.52 g KCl, 0.312 g MgSO_4_.7H_2_O, 1.52 g KH_2_PO_4_, 0.01 g Vitamin B1, 1 mL 1000× trace elements, 10 g D-glucose and 20 g agar powder in 1 L of ddH_2_O) and terrific broth 3 (TB3: 6 g casein hydrolysate, 6 g yeast extract, 200 g sucrose and 20 g agar powder in 1 L ddH_2_O). All the media used in this experiment were autoclaved for 20 min at 121°C. Autoclaved media were allowed to cool to about 50°C and dispensed into sterile 70 × 15-mm Petri plates at about 15 ml per plate. Each dish was inoculated in the center with a block of agar from the stock culture of Guy11 or the mutant strains. Unless otherwise stated, all the cultures were incubated at 26°C under diurnal fluorescent light (12/12-h light/darkness cycle). Medium treatments have three independent biological experiments with five technical replicates each time, unless otherwise stated. Colony diameters were measured after 10 days of incubation. All the pH was maintained at 7.0 unless otherwise stated.

The *Escherichia coli* strain (DH5α) was to propagate vectors. *DH5α* was grown in liquid or on solid Lysogeny broth (LB: 10 g tryptone, 5 g yeast extract and 10 g NaCl in 1 L of ddH_2_O. pH 7.0) supplemented with or without ampicillin antibiotic (Solarbio Tech. Co., Ltd, Beijing, China).

Plasmids pCX64 (for gene knockout), pKNTG-GFP (for complementation and protein expression) and RFP-HDEL (for expression and colocalization study) used in this study were sourced from the State Key Laboratory for Plant-Microbe Interaction, Plant Protection College, Fujian Agriculture and Forestry University, Fuzhou 350002, China.

### Bioinformatic analysis

For the identification of candidate components of Get proteins in *M. oryzae*, the amino acid sequences of GETs (ScGet1, ScGet2, ScGet3, ScGet4 and ScSgt2) from *Saccharomyces cerevisiae* were used as references to perform Blastp similarity search against *M. oryzae* proteome on NCBI (https://www.ncbi.nlm.nih.gov/) database accessed on 1 December 2022) and validated on FungiDB (https://fungidb.org, accessed on 2 December 2022). The identified amino sequences were subsequently named as MoGet1, MoGet2, MoGet3, MoGet4 and MoSgt2 respectively. The Get proteins of other fungi were identified the same way as the *M. oryaze* Get component proteins. SMARTdatabase (http://smart.embl-heidelberg.de) was used for domain prediction and was accessed on 30 November 2023. Finally, GPS IBS [29] and MEGA7 [30] softwares were used to construct the domain architecture and phylogenetic tree, respectively, of the proteins [12, 31].

### Generation of *M. oryzae* targeted gene deletion mutants

To generate deletion mutants, homologous recombination approach was adopted to replace the target genes (*MoGET1*, *MoGET2*, *MoGET3*, *MoGET4* and *MoSGT2*) with hygromycin-resistant fragment as previously described [12, 32, 33]. Briefly, the upstream (A fragment) and downstream (B fragment) flanking regions of the genes were amplified from *M. oryzae* genomic DNA using the primer pairs AF/AR and BF/BR, respectively. Also, HY and YG fragments of hygromycin phosphotransferase (*HPH*) gene were amplified from the pCX62 vector using primer pairs HYG-F/HY-R and YG-F/HYG-R, respectively. The A and B fragments were then fused with HY and YG divisions to obtain AH and BH fragments, respectively, by simultaneous overlap extension PCR (SOE-PCR) (Supplemental S1 Fig). All the fragments were amplified using Phanta^R^ Max Super-Fidelity DNA polymerase kit (Vazyme Biotech Co. Ltd, Nanjing, China). All primers used in this study are listed in S1 Table.

Protoplast was isolated from wild type Guy11 strain and genetic transformation was conducted using Polyethylene glycol (PEG)-mediated transformation method as previously described [34]. Candidate transformants were selected on TB3 solid media supplemented with 100 µg/mL hygromycin B or G418. Putative transformants were screened by PCR using primer pairs OF/OR and AUF/H853 or OR-F and GFP-R and the positive transformants confirmed by southern blot analysis (S1 Fig).

### Generation of complementation strains

Complementation strains for Δ*Moget1*, Δ*Moget2*, Δ*Moget3*, Δ*Moget4* and Δ*Mosgt2*, were generated by amplifying the entire ORFs sequences of *MoGET1*, *MoGET2*, *MoGET3*, *MoGET4* and *MoSGT2* and their respective native promoter from the Wt Guy11strain. Targeted bands were purified and cloned behind GFP in pKNT plasmid and constructs were transformed into the protoplasts of their respective mutant strains. The transformants were then screened by PCR and confirmed by Southern blot analyses.

To further determine the subcellular localization of MoGet1 and MoGet2 in *M. oryzae*, we constructed the GFP expression vectors pYF11: MoGet1 and pYF11::MoGet2 of MoGet1 and MoGet2, respectively, and introduced them into the wild-type strain. The transformants were screened using PCR and ascertained by laser confocal microscopy. Further, RFP-HDEL (endoplasmic reticulum Marker protein) was introduced into the respective positive transformants obtained to ascertain the organellar localization of the MoGet1 and MoGet1.

### DNA extraction, gel electrophoresis and southern blot analysis

Fungal genomic DNA was extracted from lyophilized mycelia of the Guy11, mutants and complemented strains using cetyltrimethylammonium bromide (CTAB) [31] protocol. The DNA concentration and purity were measured using Nanodrop (ThermoFisher Scientic, Basingstoke, UK). Gel electrophoresis, enzymatic DNA digestion and purification, ligation and Southern blot hybridization was conducted according to the procedures described previously [31, 34]. Probing, hybridization, staining and balance were performed using the DIG HIGH Prime DNA Labelling and Detection Starter Kit I (Roche Diagnostics GmbH, Mannheim, Germany).

### Vegetative growth, osmotic stress, cell wall integrity and reactive oxygen species sensitivity analysis

For vegetative growth assay, the WT Guy11, the GETs-deficient mutants and their corresponding complemented strains were cultured on CM, CM II, OA, SDC and MM solid media. The cultures were incubated at 26°C, 45% relative humidity (RH) in diurnal light for ten days. The diameter of each colony was measured using a meter rule in two perpendicular dimensions and the average of the two measurements was taken after subtracting the 5 mm diameter of the colonized plug.

For sensitivity assays, wild-type Guy11 strain and the mutant strains were inoculated on CM II solid media supplemented separately with osmotic stressors (NaCl, KCl and Sorbitol), cell wall and membrane stressors (CR, CFW, SDS and DTT) and different concentrations of ROS H_2_O_2_ agent; and incubated for 10 days at 26°C, 45% RH and 12 h light/12 h dark photoperiod. Colony diameters was evaluated between treated and non-treated groups.

To examine the cell wall integrity, the WT and the mutant strains Δ*Moget1*, Δ*Moget2* and Δ*Moget3* were treated with lysis enzyme following the previously established protocol [31, 34, 35] with slight modification. Briefly, 3-day old mycelia grown in CM liquid media was ground using a sterile laboratory mortar and transferred into fresh CM media. The culture was re-incubated at 26°C, 110 rpm for 12 hrs. Mycelia were filtered and 0.2 g lysing enzyme (SIGMA-ALDRICH Co., St. Louis, USA) in 20 mL 1 M C_6_H_14_O_6_ (sorbitol) added to 2 g of the wet weight mycelia. Protoplast release by each strain was estimated using haemocytomer. All the experiments were conducted in a sterile laminar flow hood. All the experiments were repeated 3 times, with 3 replicates each time.

### Conidiophore formation, conidiation and pathogenicity assay

Conidiophore development assays was performed by inoculating 5-day old mycelial plugs of Guy11, the mutants under study and their complemented strains on conidia- inducing Rice Bran agar media and incubated them at 26°C for 7 days. After, the aerial mycelium of each strain was scrubbed off and about 1 cm × 0.5 cm mycelial plug was excised and laid on a slide. The slide was incubated in 90-mm Petri plate, incubated at 27°C under humid condition, and observed under light microscope at different time points of 12 -, 24 - and 48 hrs. For conidiation assay, the cultures were re-intubated in continuous white fluorescent light for 3 days at 27°c after scrapping off of the vegetative mycelia mass to induce sporulation. Spore suspension was prepared from each strain and estimated independently under light microscope using haemocytometer.

To test the pathogenicity of each strain, an edge of growing 5-day old mycelia of each strain was excised and inoculated on 10-day old barley leaf (Gold Promise cultivar) and incubated in the dark for 24 hrs and then transferred to diurnal at 27°C for 6 days. Similarly, spore suspensions prepared from wild type control, Δ*Moget3*, Δ*Moget4* and Δ*Mosgt2* were spray - or punch inoculated on 3-week or 6-week old rice leaves. The infected seedlings were incubated in the dark for 24 hrs and diurnal for 5 days at room temperature and about 85% RH.

### *In vivo* penetration assay and life cell imaging

Host penetration and invasive hyphal growth assays were examined by inoculating 10- day old barley leaves and incubated in the dark at 26°C and under humid conditions. The leaf sheet was peeled at different time point of 12 -, 16 - and 24 hpi and examined under microscope as described previously [12, 36].

### RNA extraction for qPCR

To study the expression of hydrophobin genes in Δ*Moget1/*Δ*Moget2*, 3-day-old mycelia samples of Guy11 and Δ*Moget1/*Δ*Moget2 were* grown in CM broth at 26°C and 110 rpm. Mycelia were then filtered and washed twice using double distilled water. Total RNA was extracted from the samples using the Eastep^®^ Super RNA extraction kit (Promega Biotech Co. Ltd, Beijing, China) according to the manufacturer’s instructions. cDNA was synthesized from the total RNA by reverse transcription PCR using the Evo M-MLV RT kit with gDNA clean for qPCR (Accurate Biotechnology Co. Ltd, Hunan, China) according to the manufacturer’s instructions. Fluorescence quantitative real-time PCR was conducted using ChamQ Universal SYBR qPCR Master Mix as recommended by the manufacturer (Vazyme Biotech Co. Ltd, Nanjing, China). Three biological replicates with 3 technical replicates per biological replicate was applied to the experiments.

### Yeast two-hybrid assay

Yeast two-hybrid analysis to examine the interaction of MoGet1, -2 and -3 was performed using the MATCHMAKER GAL4 two-hybrid system 3 (Takara Bio, San Jose, USA). The protein-coding regions of the three genes were amplified from Guy11 wild-type cDNA with the primer pairs (S1 Table). MoGet1, MoGet2 and MoGet3 were cloned in the pGBKT7 bait vector, while MoGet1 or MoGet3 in the pGADT7 as prey vector. The pGBKT7 and pGADT7 were digested with *NdeI* and *EcoRI* restriction enzymes, as described previously [37]). The AD and BD constructs were cotransformed into AH109 *S. cerevisiae* strain [38]. pGBKT7-53/pGADT7-T and pGBKT7- Lam/pGADT7-T vectors were used as positive and negative controls, respectively. The emerged yeast colonies in SD/-Leu/-Trp media were isolated and cultivated on SD- four-deficient selective media (SD/-His-Leu-Trp-Ade) supplemented with 40 µg/mL X-gal for color development.

### Protein extraction, co-immunoprecipitation (Co-IP) and western blot assay

Total protein was extracted from 3-day old *M. oryzae* mycelia of Moget1-RFP, MoGet3-RFP, MoGet1-GFP and GFP-MoGet2 according to previous protocol [39]. Briefly, about 5 g mycelial powder prepared by grinding lyophilized 3-day old mycelia was added into a sterile 2-mL Eppendorf tube. 1 mL Lysis buffer (10 mM Tris/Cl pH=7.5, 150 mM NaCl, 0.5 mM EDTA, 0.5% NP-40), 10 μL PMSF, 10 μL protein inhibitor in the ratio of 100:1:1 were added. The suspension was vortexed to homogenize, incubated in ice for 10 min and centrifuged at 4°C, 14000 rpm for 15 min (Micro-centrifuge 5430 R). The supernate was collected into a 1.5-ml Eppendorf centrifuge tube and 5x SDS buffer was added at the ratio of 5mL supernate to 1 ul 5x SDS buffer. The sample was boiled for 10 min for denaturing and stored at – 20°C for western blot analysis.

For Co-IP assay, GFP-fusion proteins were isolated and incubated with 30 ul GFP-Trap magnetic beads (ChromoTek, Martinsried, Germany) according to manufacturer’s instructions. To obtain the protein from GFP-Trap magnetic beads, the sample-containing 10-mL tubes were placed in ice for 10 min for magnetic beads sedimentation. Then, the tubes were transferred to a magnetic rack, allowed for 1 min to trap the GFP-bound beads and the supernate was carefully discarded. The beads were washed three times with dilution buffer (50 mM Tris, 150 mM NaCl, pH 7.4) and then elusion buffer. Equal volume of protein loading buffer was added, then denatured by boiling and placed on a magnetic frame for 1 min. The elusion protein was collected and 40 ul of total protein was loaded in 10% SDS-PAGE gel for immunoblotting and co- immunoprecipitation analysis.

For phosphorylation signal, total protein from the mycelial of Guy11 and Δ*Moget1/*Δ*Moget2* double mutant was extracted as previously described [40]. The phosphorylation level of Mps1 was detected through western blot analysis. Briefly, 40 µl of total protein sample was loaded into each 10% SDS-PAGE gel well and separated by electrophoresis. The gel was transferred onto a nitrocellulose membrane (Amersham, Piscataway, NJ, USA) for blotting. The membrane was incubated in p44/42 MAPK antibody (Cell Signaling Technology, Beverly, MA, USA). Chemiluminescent signals of the specific protein bands were detected using ECL kit (Amersham Biosciences, Freiburg, Germany).

### Microscopic observation

Hyphal penetration and invasion on host media surfaces, GFP and RFP localizations were observed under a laser confocal microscope equipped with a Nikon A1 Plus imaging instrument (Tokyo, Japan).

### Statistical analysis and reproducibility

Quantification of growth diameter, sensitivity of the strains to stressors, conidiation and lesion types was performed using Microsoft excel spreadsheet and GraphPad prism5(cite). ImageJ software, Microsoft excel spreadsheet and GraphPad prism5 were used to conduct disease area quantification. Error bars represent standard deviation from the mean in all the Figures, and *p* values were determined by one- or two-way ANOVA. All experiments were repeated as stated in each Figure legends.

## Results

### Identification of Get proteins in *M. oryzae*

Amino acid sequences of Get1, Get2, Get3, Get4 and Sgt2 from *Saccharomyces cerevisiae* (S288C) were used to perform a BLASTp search at the FungiDB (https://fungidb.org, accessed on 30 October 2022) and National Centre for Biotechnology Information (https://www.ncbi.nlm.nih.gov/, accessed on 30 October 2022). The identified homologs were named MoGet1, MoGet2, MoGet3, MoGet4 and MoSgt2, respectively. Compare to *S. cerevisiae* Get components, alignment results showed no significant similarity for MoGet1 and MoGet2 (Fig 1A and 1B), but 51.30%, 33.55% and 34.01% sequence similarities were obtained for MoGet3, MoGet4 and MoSgt2, respectively. More homologs of these proteins were found in the other eight top important pathogenic fungi [41], and further analyzed at SMARTdatabase (http://smart.embl-heidelberg.de/, accessed on 14 October 2023). Protein domain prediction showed that MoGet1 and MoGet2 contain transmembrane domain (TMD), unlike cytosolic MoGet3 and MoGet4 that contain only low complexity regions (LCRs) [42, 43]. Except for Get1 of *Fusarium graminaerum*, *Blumeria graminis*, *Colletorichum siamense* and *S. cerevisiae* and Get2 of *Blumeria graminis, Uslago maydis* and *Homo sapiens*, TMD insertase is conserved in all fungi and mammals analyzed (Fig 1A and 1B). On the other hand, MoSgt2 possesses a TPR catalytic domain, which is conserved in the top nine important plant pathogenic fungi as well as in *N. crasa*, *S. cerevisiae* and *H. sapiens* (Fig 1E). Moreover, the phylogenetic analysis indicates that MoGet1, MoGet2, MoGet3, MoGet4 and MoSgt2 have close ancestry with Gets and Sgt2 in all the fungi analyzed, except that they have distant ancestry with ScGet1, HsGet2, HsGet3, HsGet4 and HsSgt2, respectively (Fig 1F - 1J).

**Fig 1.**
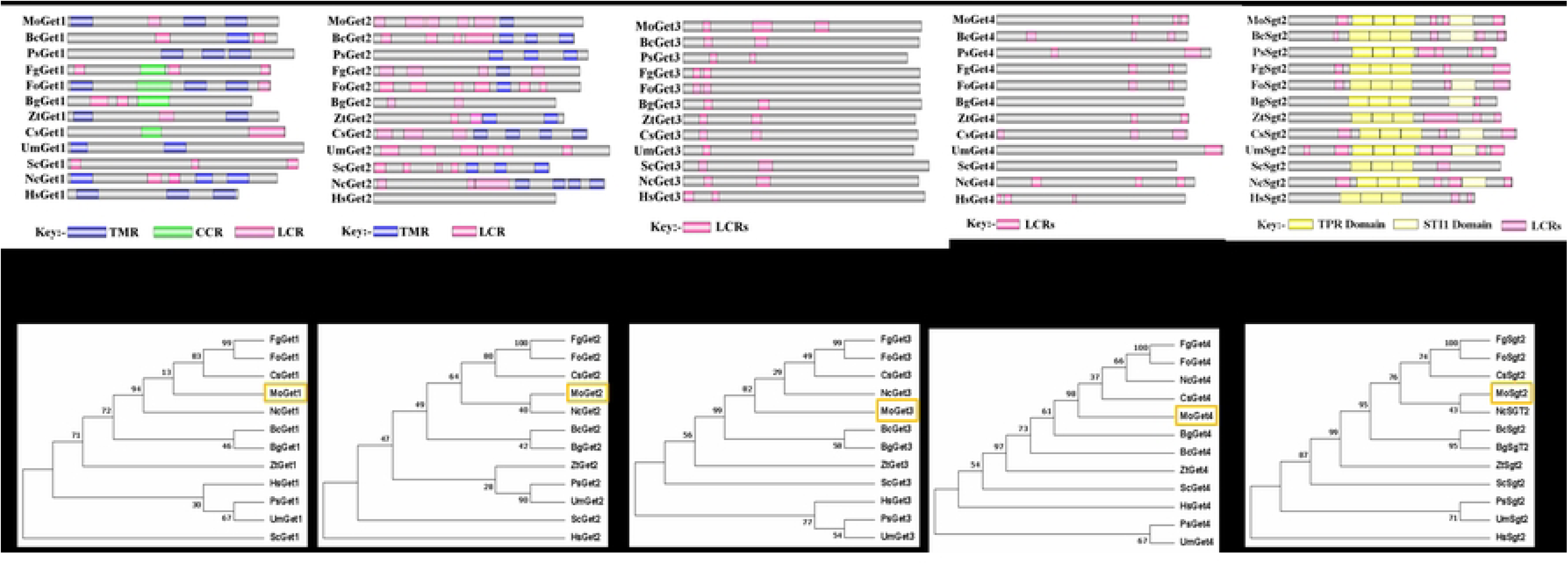
Domain architecture and phylogeny of Gets in phytopathogenic fungi, *Neurospora crasa* and *Homo sapiens*: *S. cerevisiae* Get1 (NP_011495.1), ScGet2 (NP_011006.2), ScGet3 (NP_010183.1), ScGet4 (NP_014807.3), ScSgt2 (NP_014649.1); *M. oryzae* Get1 (XP_003715385.1), MoGet2 (XP_003717841.1), MoGet3 (XP_003712205.1), MoGet4 (XP_003715022.1), MoSgt2 (XP_003710298.1); *Botrytis cinerea* Get1 (XP_024548203.1), BcGet2 (EMR80383.1), BcGet3 (XP_001552593.1), BcGet4 (XP_024553867.1), BcSgt2 (XP_001556769.1); *Puccinia striiformis* Get1 (KAI7967766.1), PsGet2 (XP_047806390.1), PsGet3 (KAI7934910.1), PsGet4 (KNE97063.1), PsSgt2 (XP_047807857.1); *Fusarium graminearum* Get1 (XP_011321211.1), FgGet2 (QPC79602.1), FgGet3 (XP_011318797.1), FgGet4 (XP_011327068.1), FgSgt2 (XP_011321549.1); *F. oxysporum* Get1 (KAG7407779.1), FoGet2 (TXC07091.1), FoGet3 (XP_018240895.1), FoGet4 (XP_018234399.1), FoSgt2 (KAH7214056.1); *Blumeria graminis* Get1 (EPQ67104.1), BgGet2 (CAD6504456.1), BgGet3 (EPQ64508.1), BgGet4 (EPQ61584.1), BgSgt2 (EPQ62108.1); *Zymoseptoria tritici* Get1 (XP_003848188.1), ZtGet2 (XP_003856636.1), ZtGet3 (XP_003853017.1), ZtGet4 (XP_003850056.1), ZtSgt2 (XP_003857067.1); *Colletotrichum fructicola* Get1 (XP_036499662.1), CfGet2 (XP_036499714.1), CfGet3 (XP_036489687.1), CfGet4 (XP_036498048.1), CfSgt2 (XP_036493300.1); *Ustilago maydis* Get1 (XP_011386800.1), UmGet2 (XP_011391811.1), UmGet3 (XP_011390272.1), UmGet4 (XP_011387570.1), UmSgt2 (XP_011392129.1); *Neurospora crasa* Get1 (XP_957971.1), NcGet2 (XP_963518.1), NcGet3 (XP_960897.1), NcGet4 (XP_961556.1), NcSgt2 (XP_001728325.1) and Homo sapiens Get1 (NP_004618.2), HsGet2 (NP_001736.1), HsGet3 (NP_004308.2), HsGet4 (NP_057033.2) and HsSgt2 (NP_061945.1). (A and F) Get1 domain architecture and phylogenetic analysis in different fungi. (B and G) Get2 domain architecture and phylogenetic analysis in different fungi. (C and H) Get3 domain architecture and phylogenetic analysis in different fungi. (D and I) Get4 domain architecture and phylogenetic analysis in different fungi. (Fig E and J) Sgt2 domain architecture and phylogenetic analysis in different fungi. The evolutionary history was inferred using the maximum likelihood method based on the JTT matrix-based model. The analysis involved 60 amino acid sequences. The maximum likelihood phylogeny for the amino acids was tested with 1000 bootstrap replicates. Evolutionary analyses were conducted using MEGA7. TMR: Transmembrane region; CCR: Coil-coil region; TPR: Tetratricopeptide repeat; STI1: Stress inducible 1; LCRs: low complexity regions.

### MoGet1 and MoGet2 are crucial for vegetative growth and conidiation

To determine the role of MoGet1, MoGet2, MoGet3, MoGet4 and MoSgt2 in vegetative growth of *M. oryzae*, the *MoGET1*-, *MoGET2*-, *MoGET3*-, *MoGET4*- and *MoSGT2*- deficient strains alongside with WT type Guy11 and the complementation were cultured on CM, CM II, OA, SDC and MM solid media at 26°C and colony diameters measured after 10 days. Our results showed that the growths of Δ*Moget1* and Δ*Moget2* mutants were significantly reduced on all the growth media used compared to Guy11 and their respective complemented strains (Fig 2A, 2B and Table 1). The Δ*Moget1* mutant exhibited a more redistricted colony expansion, while the Δ*MoGet2* produced thin aerial hyphae (Fig 2C). Conversely, Δ*Moget3* mutant showed significant increase in vegetative growth compared to the Guy11 wild-type and complemented strains (Fig 2A, 2B and Table 1), demonstrating that MoGet3 negatively regulates the fungal vegetative growth. On the other hand, the *MoGET4* and *MoSGT2* gene deletion mutants are similar to Guy11 in vegetative growth. Reintroduction of the *MoGET1, MoGET2*, *MoGET3*, *MoGET4* and *MoSGT2* genes into the mutant strains restored their normal growths, except for the Δ*Moget2*/*MoGET2* complemented strain which is restricted in growth and sporulation, and Δ*Moget3/MoGET3*, which outperformed Guy11 in hyphal growth (Fig 2A and 2B). These results demonstrate that MoGet1 and MoGet2 are important for normal vegetative growth, while MoGet3 negatively regulates mycelial development in *M. oryzae*.

**Fig 2.**
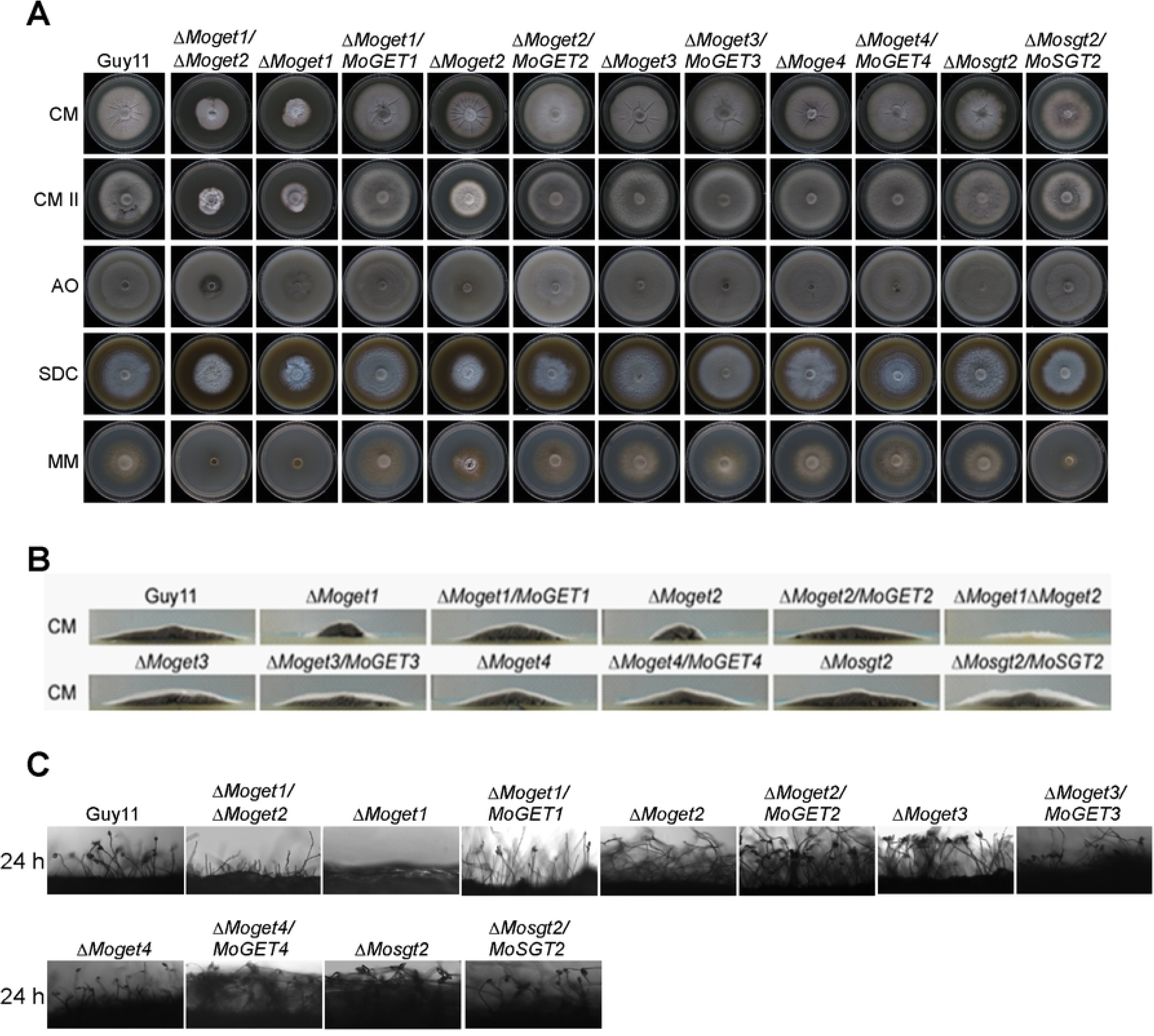
Role of MoGet1, MoGet2, MoGet3, MoGet4 and MoSgt2 in hyphal growth in *M. oryzae*. (A and B) Vegetative growth of the Guy11, mutants and their complemented strains on CM, CM II, OA, SDC and MM media at 10 days post inoculation (dpi). (C) Conidiophores and conidia formation of the conidiophores of Guy11, the mutants and complemented strains on rice bran medium.

**Table 1.** Comparison of growths and conidiation of the mutant strains.

| Strain | Colony diameter (cm)<sup>g</sup> |  |  |  |  | Conidiation<sup>s</sup><br>(×10<sup>4</sup>) |
| --- | --- | --- | --- | --- | --- | --- |
|  | CM | CM II | OA | SDC | MM |  |
| Guy11 | 4.9±0.33<sup>b</sup> | 5.4±0.18<sup>b</sup> | 5.1±0.23<sup>b</sup> | 5.3±0.16<sup>b</sup> | 4.6±0.65<sup>a</sup> | 202.92±72.30<sup>b</sup> |
| ΔMoget1/<br>ΔMoget2 | 3.3±0.49<sup>d</sup> | 3.3±0.64<sup>d</sup> | 3.1±0.33<sup>d</sup> | 3.1±0.56<sup>e</sup> | 2.8±0.83<sup>c</sup> | 0.00±0.00<sup>RO</sup> |
| ΔMoget1 | 2.1±0.63<sup>d</sup> | 2.3±0.21<sup>e</sup> | 2.7±0.59<sup>e</sup> | 2.2±0.79<sup>f</sup> | 2.0±0.52<sup>d</sup> | 0.00±0.00<sup>RO</sup> |
| ΔMoget1/<br>MoGET1 | 5.0±0.22<sup>b</sup> | 4.6±1.11<sup>c</sup> | 5.1±0.17<sup>b</sup> | 4.9±0.52<sup>b</sup> | 4.9±0.40<sup>a</sup> | 141.58±54.79<sup>c</sup> |
| ΔMoget2 | 3.9±0.38<sup>c</sup> | 3.9±0.49<sup>c</sup> | 4.2±0.31<sup>c</sup> | 3.9±0.22<sup>d</sup> | 3.5±0.31<sup>b</sup> | 0.00±0.00<sup>RO</sup> |
| ΔMoget2/<br>MoGET2 | 4.9±0.74<sup>b</sup> | 4.6±0.40<sup>c</sup> | 4.8±0.32<sup>b</sup> | 5.0±0.69<sup>b</sup> | 4.7±0.50<sup>a</sup> | 105.75±13.30<sup>d</sup> |
| ΔMoget3 | 5.6±0.24<sup>a</sup> | 5.8±0.13<sup>a</sup> | 5.9±0.095<sup>a</sup> | 5.9±0.11<sup>a</sup> | 5.2±0.31<sup>a</sup> | 251.58±15.29<sup>a</sup> |
| ΔMoget3/<br>MoGET3 | 5.7±0.37<sup>a</sup> | 6.1±0.11<sup>a</sup> | 6.0±0.04<sup>a</sup> | 6.0±0.18<sup>a</sup> | 5.2±0.44<sup>a</sup> | 0.83±0.63<sup>f</sup> |
| ΔMoget4 | 5.3±0.45<sup>b</sup> | 5.3±0.48<sup>b</sup> | 5.8±0.13<sup>a</sup> | 5.9±0.11<sup>a</sup> | 4.8±0.58<sup>a</sup> | 200.04±21.79<sup>b</sup> |
| ΔMoget/<br>MoGET4 | 5.3±0.48<sup>b</sup> | 5.4±0.30<sup>b</sup> | 5.8±0.15<sup>a</sup> | 5.8±0.11<sup>a</sup> | 5.0±0.53<sup>a</sup> | 75.50±31.40<sup>e</sup> |
| ΔMosgt2 | 5.1±0.26<sup>b</sup> | 5.2±0.24<sup>b</sup> | 5.7±0.083<sup>a</sup> | 5.6±0.26<sup>a</sup> | 4.8±0.3<sup>a</sup> | 207.52±34.83<sup>b</sup> |
| ΔMosgt2/<br>MoSGT2 | 4.5±0.52<sup>c</sup> | 4.7±0.40<sup>c</sup> | 5.1±0.33<sup>b</sup> | 4.9±0.40<sup>b</sup> | 3.2±1.12<sup>b</sup> | 115.00±13.00<sup>d</sup> |

<sup>g</sup>Colony diameter (cm) of Guy11, the mutants and their complemented strains on CM, CM II, OA, SDC and MM at 27°C 10 dpi.
<sup>s</sup>Statistical analysis of average conidiation of the respective mutants and their complemented strains relative to Guy11 wild-type strain 10 dpi on RB medium at 27°C.
<sup>RO</sup>Rarely observed.

Conidia facilitate the survival, efficient dissemination and disease perpetuation of the fungus [44]. To unveil the role of the Get proteins in asexual sporulation of *M*. *oryzae*, we harvested, quantified and compared the amount of conidia produced by the various strains. The Δ*Moget1*, Δ*Moget2* and Δ*Moget1/*Δ*Moget2* mutant strains failed to produce spores while Δ*Moget4* and Δ*Mosgt2* strains produced similar number of spores as the wild-type strain (Table 1), suggesting that MoGet1 and MoGet2 are essential for conidiation in *M. oryzae*. Conidiophore formation is a prerequisite for conidiation and conidia-mediated infection of host plants under favourable conditions [44]. Δ*Moget1*, Δ*Moget2* and Δ*Moget1/Moget2* mutants failed to produce conidiophore (Fig 2D), which further supports their critical involvement in asexual reproduction in *M. oryzae*.

Taken together, we conclude that the *MoGET1* and *MoGET2* genes are essential for growth, development and asexual reproduction in *M. oryzae*.

Mean and standard deviation were determined using contingency table analysis with row means in Microsoft Excel spreadsheets and GraphPad Prism 5. Similar values were obtained from three independent experimental repeats with five technical replicates for each repetition.

The superscript a, b, c, d and e indicate significant changes compare to Guy11. Same letters in a column shows no significance difference.

### The role of the Get components in stress tolerance

Environmental stimuli trigger adaptive cellular responses to optimize tolerance, survival and proliferation [45, 46], and an organism’s response to stress involves the integrated function of many components of cell metabolism [46].To determine the Role of the GET components in stress response in rice blast fungus, we cultured the Guy11, Δ*Moget1*, Δ*Moget2*, Δ*Moget3*, Δ*Moget4* and Δ*Mosgt2*, and their respective complemented strains on CM II supplemented with 1 M NaCl, KCl and C_6_H_14_O_6_ (Sorbitol) (as osmotic stress-inducing agents) to evaluate their growth responses. It is obvious from the results that the growth of Δ*Moget2* mutant strain was more inhibited on media supplemented with NaCl, KCl and C_6_H_14_O_6_ than Δ*Moget1*, Δ*MoGet3*, Δ*MoGet4* and Δ*MoSgt2* strains (Fig 3 and Table 2). Interestingly, the Δ*Moget1/*Δ*Moget2* double knock mutant was less inhibited than Δ*Moget2*, suggesting that MoGet1 serves as a negative osmo-regulator in *MoGET2*-deficient mutant strain. In sorbitol-supplemented media, Δ*Moget1* was the least inhibited (P< 0.001) strain while Δ*Moget1*/Δ*Moget2*, Δ*Moget3* and all the complemented strains showed no significant difference in their growths compared to Guy11. Also, Δ*Moget4,* Δ*Mosgt2* and their complemented strains showed no significant inhibition rate as compared to Guy11 (Fig 3 and Table 2).

**Fig 3.**
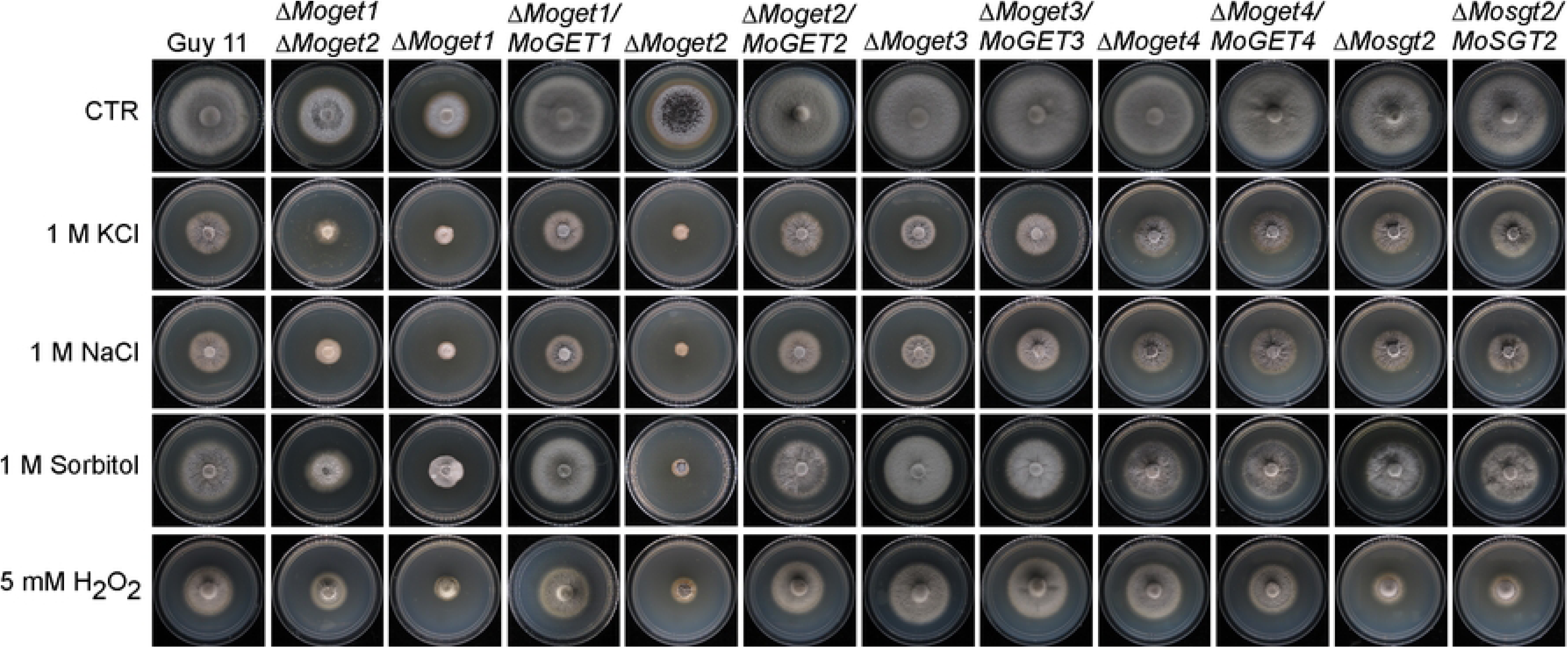
Growth phenotype and Sensitivity of MoGETs-deficient mutants to KCl, NaCl, Sorbitol and different concentrations of H_2_O_2_. Guy11 wild-type, Δ*Moget1/*Δ*Moget2*, Δ*Moget1*, Δ*Moget2*, Δ*Moget3*, Δ*Moget4*, Δ*Mosgt2* and their complemented strains were cultured on CM II supplemented with 1 M Sorbitol, KCl, NaCl and 5.0 mM H_2_O_2_, incubated at 26°C for 10 days and then sampled for sensitivity assay for osmotic and ROS stress response. The inhibition rate of each treatment was compared with the growth rate of the untreated control.

**Table 2.**
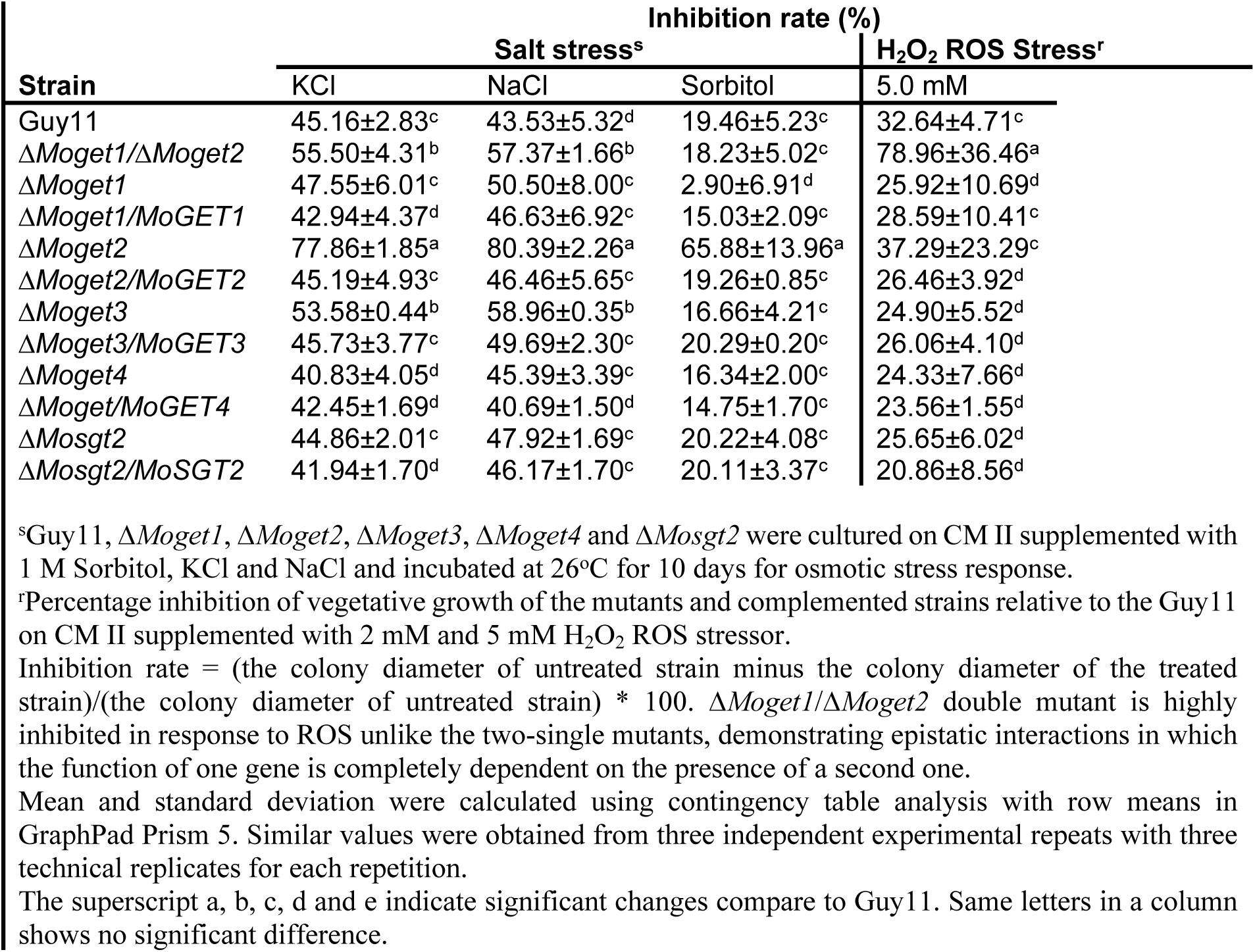
Salt and ROS stress assays of MoGets component mutants.

Reactive oxygen species (ROS) sensing and metabolism are controlled by a variety of proteins involved in oxidation-reduction reactions [47]. Therefore, we investigated the role of *M. oryzae* Get proteins in oxidative stress response by growing the various strains on CM II medium supplemented with 2.5 mM or 5 mM H_2_O_2_ at 26°C for 10 days (Paul D. Ray et al., 2012; Biregeya, J., et al., 20). The results revealed that the Δ*Moget1/*Δ*Moget2* double mutant was significantly (*** p < 0.001) inhibited on media containing 5 mM H_2_O_2_ compared to the single mutants, their complemented strains and Guy11 (Fig 3, Table 2), suggesting a functional redundancy of the two proteins during oxidative stress response. On the other hand, no significant difference was observed among Δ*Moget3*, Δ*Moget4* and Δ*Mosgt2* in response to the H_2_O_2_-induced stress, compared to Guy11.

Cell wall maintains cell morphology, protects the cell contents and mediates the transmission of external stimuli into the cell [48]. To establish the contribution of the GET genes to cell wall and cell membrane integrity in *M. oryzae*, we cultured the various fungal strains Δ*Moget1*, Δ*Moget2* Δ*Moget1*/Δ*Moget2,* Δ*MoGet3*, Δ*MoGet4* and Δ*MoSgt2* on CM II media supplemented with the cell wall and cell membrane stressors calcofluor white (CFW), Congo red (CR), sodium dodecyl sulphate (SDS) And dithiothreitol (DDT). After 10 days of incubation at 26°C, we analysed the colony diameter of each strain. Results showed that the growth of Δ*Moget3* is less inhibited by CR and CFW than Guy11 (Fig 4A and Table 3), suggesting that MoGet3 is also a negative regulator of cell wall integrity. The Δ*Moget2* mutant, on the other hand, is highly inhibited by CFW, an indication that MoGet2 is required for cell wall integrity in M. oryzae. Meanwhile, Δ*Moget1*/Δ*Moget2* double mutant was less inhibited by CFW compared to ΔMoget2 after a 10-day incubation. This result indicates that MoGet1 is necessary for the ΔMoget2-induced cell wall compromise. On DDT-supplemented CM II media, our data showed that Δ*Moget1* and Δ*Moget2* are significantly inhibited compared to Guy11. This demonstrates that ER is compromised and MoGet1 and MoGet2 are required for ER membrane integrity.

**Fig 4.**
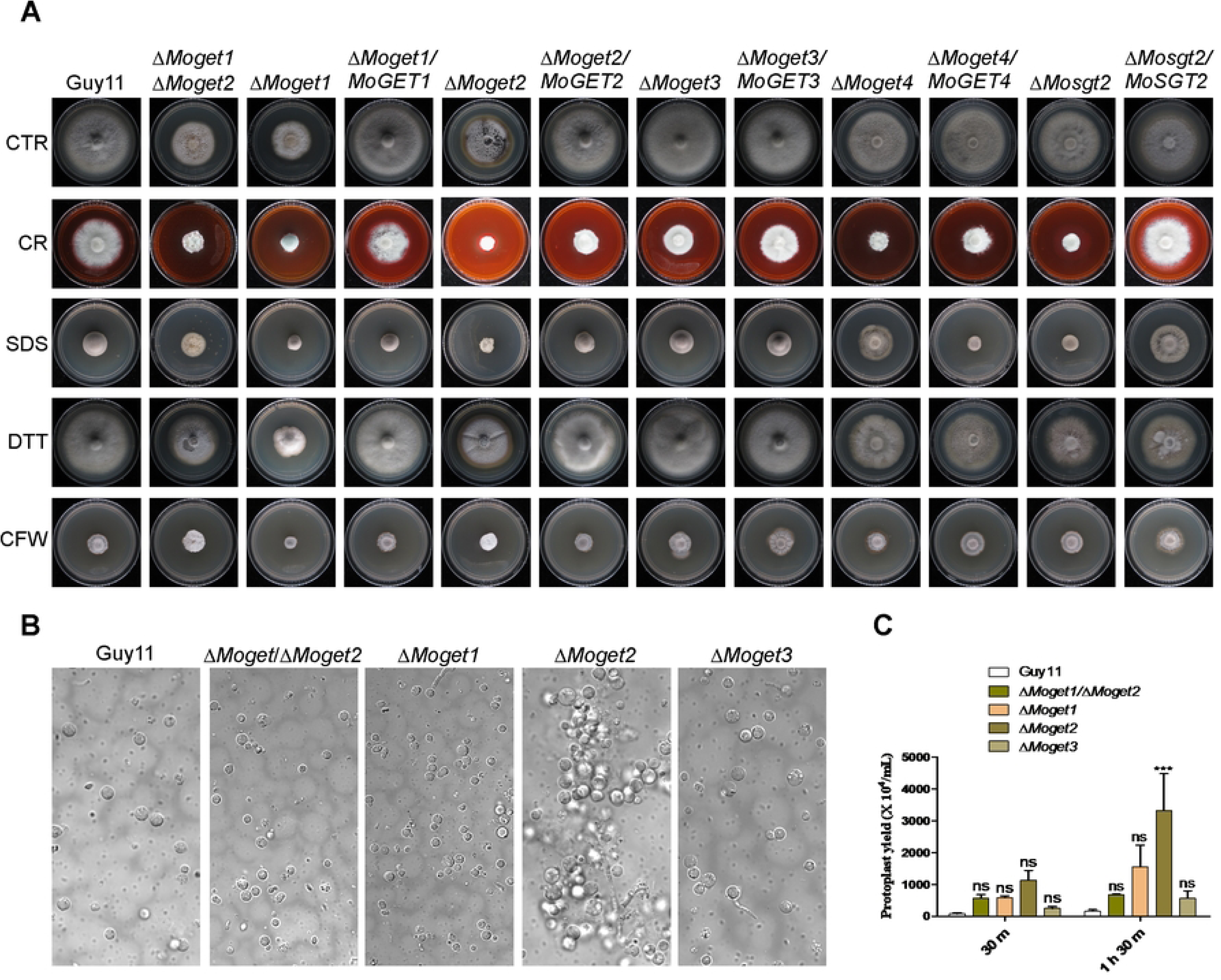
Sensitivity of Guy11, Δ*Moget1/*Δ*Moget2*, Δ*Moget1*, Δ*Moget2*, Δ*Moget3*, Δ*Moget4* and Δ*Mosgt2* to cell wall stressors. (A) Vegetative growth of the Guy11, the mutants and complemented strains cultured on CM II supplemented with the cell wall stressors (200 g/mL CR, 0.01% SDS and 200 g/mL CFW) and imaged 10 dpi. (B) Protoplasts from Guy11, Δ*Moget1/*Δ*Moget2,* Δ*Moget1*, Δ*Moget2* and Δ*Moget3* after treatment with lysing enzyme at 30°C, 85rpm at different time points: 30 m and 1 h 30 m. Scale bar =10 µm. (C) Amount of protoplast released by the Guy11, Δ*Moget1/*Δ*Moget2* double mutant and the three single mutant strains. Statistical analysis was conducted using two-way ANOVA with Bonferroni posttests (GraphPad Prism 5). Each sample was compared with Guy11 wild-type. ‘ns’ and ‘***’ represent significant differences p < 0.05 and p < 0.001, respectively. Error bar represents standard deviation from the mean. The experiments were conducted three times with five independent replicates.

**Table 3.**
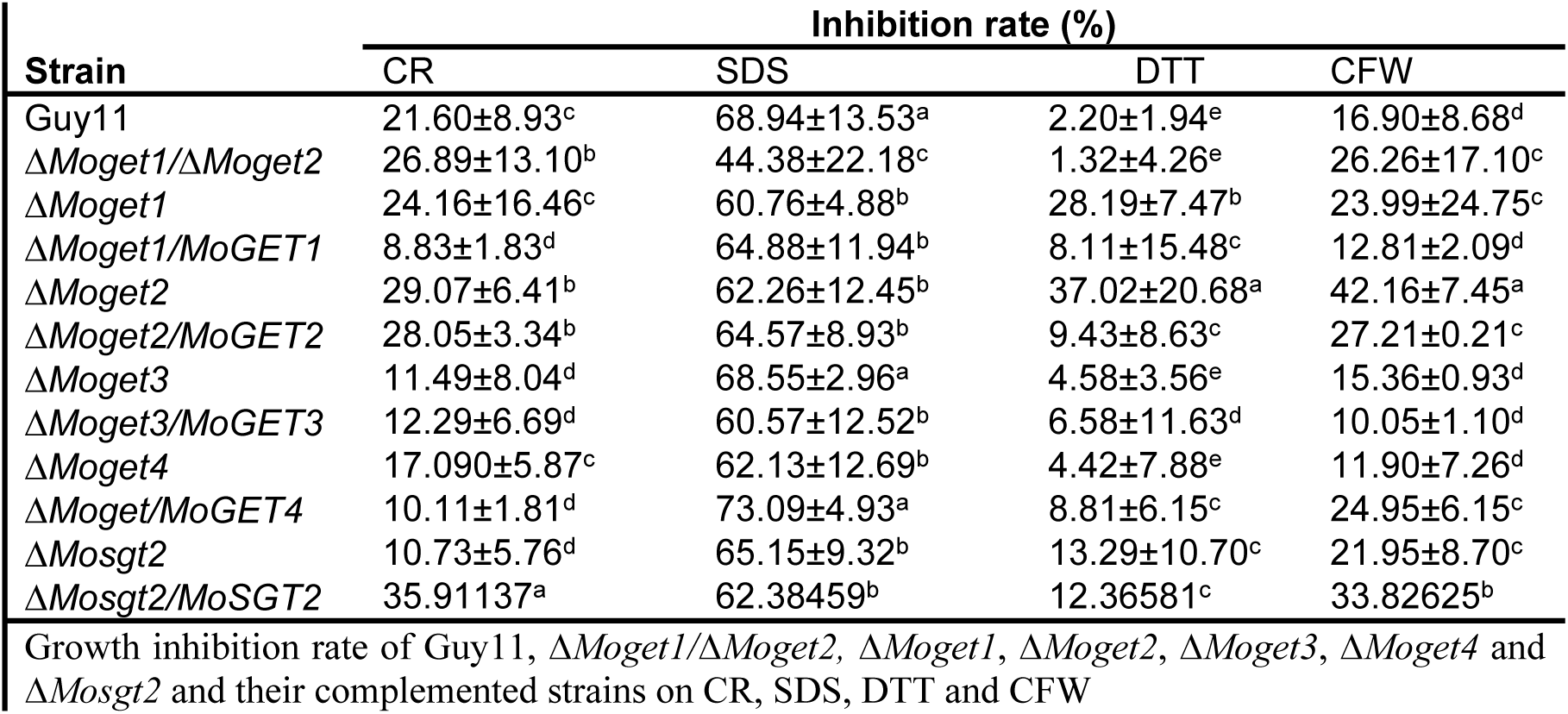
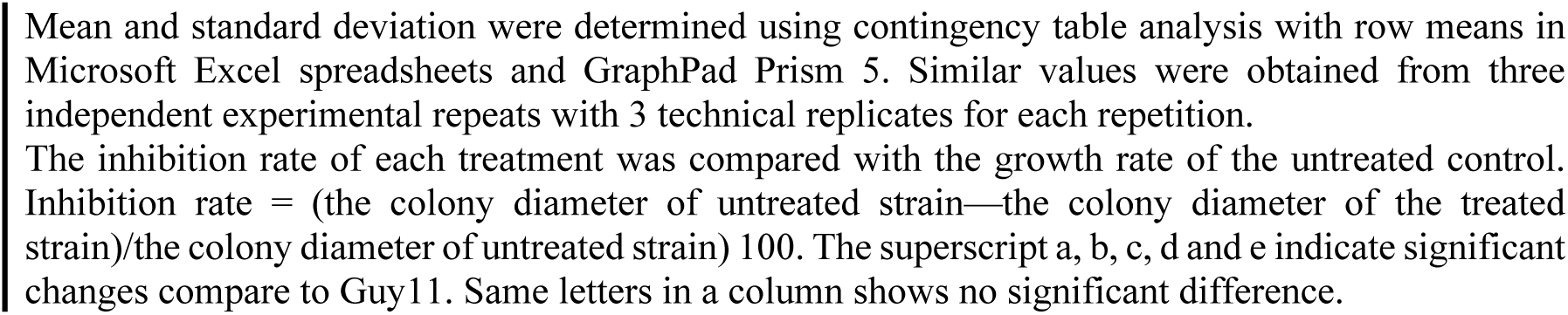
Cell wall and membrane integrity stress assay of MoGets component mutants.

To further confirm the role of MoGet2 in cell wall maintenance, we exposed the mycelia of Guy11, Δ*Moget1*/Δ*Moget2*, Δ*Moget1*, Δ*Moget*2 and Δ*Moget*3 strains to a cell wall-degrading enzyme (25 mg/mL lysing enzyme), cultured under agitation at 85 rpm, 30°C for 30 min or 1 h 30 min periods. We observed that the hyphae from the Δ*Moget2* mutant were well digested and released the highest number of protoplasts at each time point, followed by Δ*Moget1* (at 1 h 30 min), the Guy11 wild type control and then Δ*Moget3* mutant (Fig 4B and 4C), indicating that the cell wall of the Δ*Moget2* mutant was more prone to degradation than those of the other strains.

### MoGet1 and MoGet2 are critical for *M. oryzae* pathogenesis

To investigate the roles of the GET genes in the pathogenicity of *M. oryzae*, we inoculated 5-day old mycelial plug from Δ*Moget1*, Δ*Moget2*, Δ*Moget3*, Δ*Moget4* and Δ*Mosgt1* mutants on 10-day old barley leaves. The infected leaves were observed and photographed 5 days after infection. The results indicate that Δ*Moget1*, Δ*Moget2* and Δ*Moget1/*Δ*Moget2* mutants failed to cause noticeable disease symptoms on the susceptible barley leaves (Fig 5A and 5B). However, Δ*Moget3* was significantly more pathogenic than Guy11 and the other strains, suggesting that MoGet3 negatively regulates pathogenicity of *M. oryzae*.

**Fig 5.**
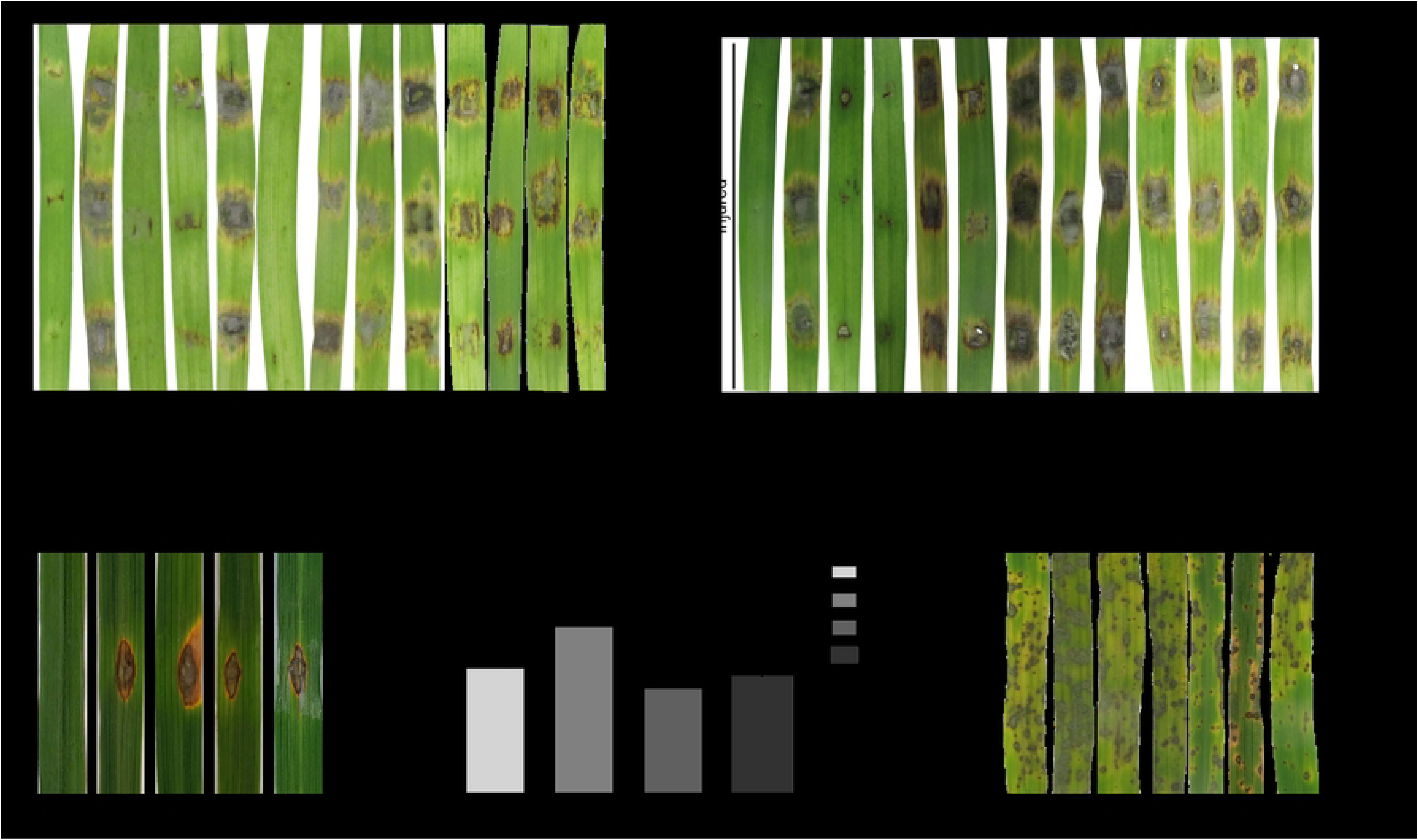
MoGet1 and MoGet2 are essential for *virulence* of *M. oryzae,* Δ*Moget3* more pathogenic than Guy11. (A and B) Disease lesions of Guy11, Δ*Moget1*, Δ*Moget2*, Δ*Moget3*, Δ*Moget4*, Δ*Mosgt2* and their complemented strains on intact and injured barley leaves, respectively. 10-day old detached barley leaves were wounded (or left intact) with a pipette tip, inoculated with 5-mm mycelial plugs, incubated in the dark for 24 h and then transferred to fluorescence continuous light for 4 days at 27°C. (C) Disease lesions of Guy11 and spore-producing mutants of Δ*Moget3*, Δ*Moget4* and Δ*Mosgt2* on host rice leaves. 6-week old Rice leaves were punch-inoculated, kept in the dark for 24 h and transferred to 12 h/12 h diurnal light under humid conditions. Photograph was taken 14 dpi. (D) Statistical analysis of Percentage disease area of representative leaves of Guy11, Δ*Moget3*, Δ*Moget4* and Δ*Mosgt2*. ImageJ software and one-way ANOVA with Tukey’s multiple-comparison test (GraphPad Prism 5) were used to perform statistical analysis. (E) Spray inoculation on 3-weeks old host rice leaves using spore suspension prepared from Guy11, Δ*Moget3*, Δ*Moget4* and Δ*Mosgt2.* The inoculated 3-week old rice seedling were maintained in the humid and dark condition for 24 hours and transferred to 12 h/12 h diurnal light. Photograph was taken 5 dpi. Asterisk = p-value (*, p < 0.05). Error bar represents the standard deviation from the mean of three independent repeats.

For conidia-mediated infection, we punch-inoculated 6-week old or spray-inoculated 22-day old seedlings of the susceptible rice cultivar CO-39 with spore suspensions from Guy11 and the various conidia-producing mutants (Δ*Moget1*, Δ*Moget2* and Δ*Moget1/*Δ*Moget2* do not produce conidia) and assessed the disease symptoms after 10 d for punch inoculation and 5 d for spray inoculation. For punch inoculation, all the strains were able to cause infection, with Δ*Moget3* displaying more expanded disease area (Fig 5C and 5D). Similarly, results of the spray inoculation showed that both Δ*Moget4* and Δ*Mosgt2* exhibited similar virulence, while Δ*Moget3* was more virulent (Fig 5E) compare to Guy11.

### Appressorium formation and penetration assay of the Get mutants

Appressorium formation is necessary for host penetration and colonization by *M. oryzae*. Previous studies have shown that hyphal-mediated infection of the host leaf by *M. oryzae* involves the pre-formation of an appressorium-like structure at the tip of the hyphae [12, 44]. We reasoned out that the inability of Δ*Moget1*, Δ*Moget2* and Δ*Moget1/*Δ*Moget2* mutants to cause infection might be due to their failure to form appressoria. To test this, we inoculated mycelial plugs from 5-day old cultures of the Guy11, the non-spore producing mutants Δ*Moget1*, Δ*Moget2* and Δ*Moget1*/Δ*Moget2* on 10-day old barley (Golden Promise) leaves as well as on artificial hydrophobic cover slips, and incubated them in the dark for 24 hours. Unlike Guy11 and Δ*Moget1* mutant which produced appressoria on both natural and artificial hydrophobic surfaces, the Δ*Moget1/*Δ*Moget2* double mutant failed to form any appressorium (Fig 6A). Few appressoria were formed on barley leaf surface, but not on artificial hydrophobic surface, when inoculated with Δ*Moget2* mutant mycelia plug (Fig 6A and 6C), suggesting that MoGet1/MoGet2 play important synergistic role in appressorium formation. We also observed from the results that Δ*Moget1* and Δ*Moget2* mutants were able to penetrate the host leaf but failed to invade and cause disease symptoms after 48 to several hours of inoculation (Fig 6A).

**Fig 6.**
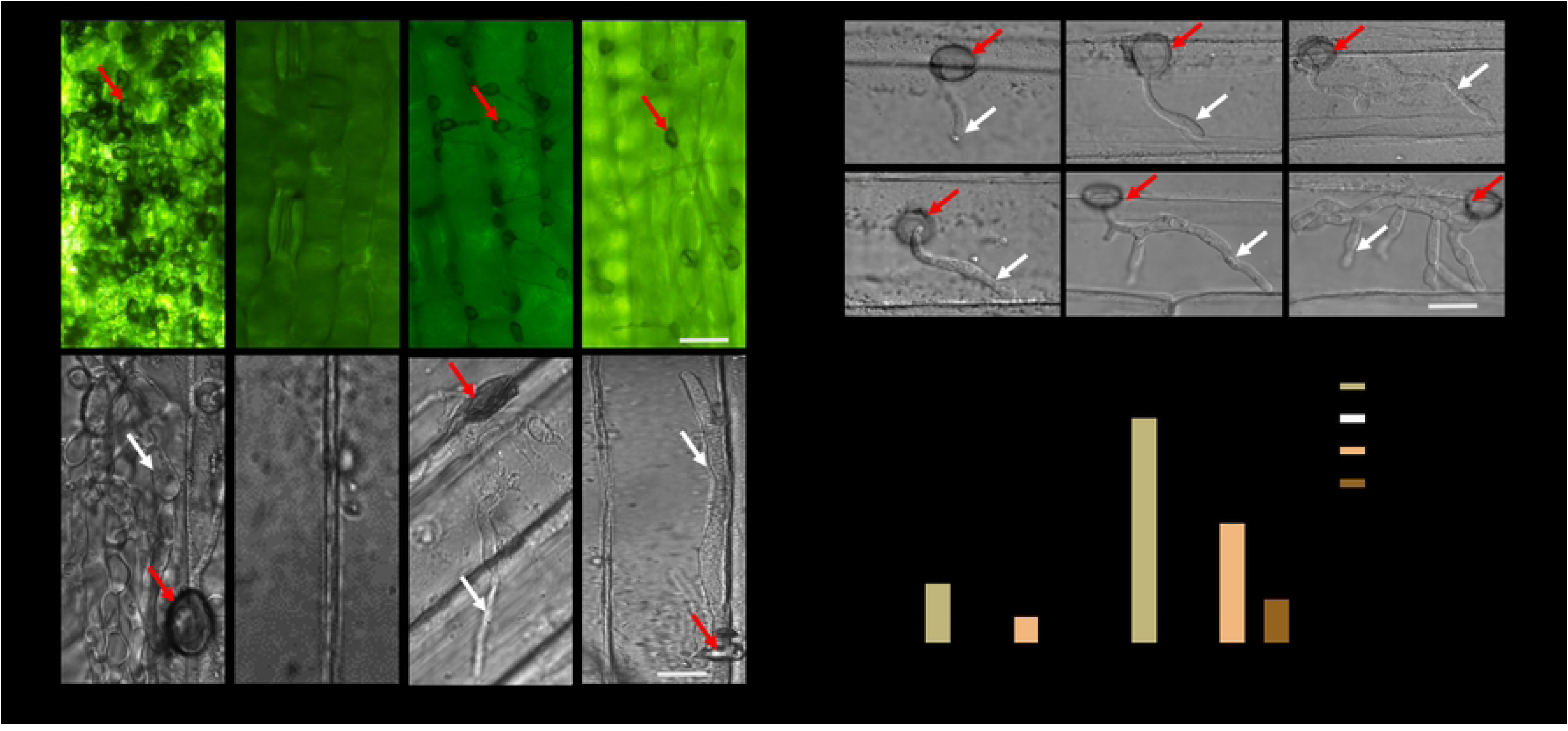
Appressorium formation and penetration assay of Guy11 and the mutant strains. (A) Hyphal-mediated appressorium formation of Guy11 wild-type, Δ*Moget1/*Δ*Moget2*, Δ*Moget1* and Δ*Moget2* mutant strains on hydrophobic barley surface and their invasion of barley epidermal cell (B) Appressorium-mediated penetration of Guy11 and Δ*Moget3* mutant strains into epidermal cell of barley leaves at 12, 16 and 24 hpi. (C) Quantification of hyphal-mediated appressorium formation on artificial and natural hydrophobic surfaces. Error bar represents the standard deviation (SD) from the mean of two independent repeats with two technical replicates. Statistical analysis was conducted using Two-way ANOVA with Bonferroni post-test (GraphPad Prism 5). Each sample was compared with Guy11 wild-type. ‘**’ and ‘***’ represent significant differences p < 0.01 and p < 0.001, respectively. AHPS: Artificial hydrophobic surface; NHPS: Natural hydrophobic surface; Red arrow: Appressorium; white arrow: Hyphae. Scale bar = 10 µm.

To establish the reason for increased pathogenicity exhibited by Δ*Moget3* mutant, detached barley leaves were inoculated with conidia suspensions (1 × 10^4^ spores/mL) from Guy11, Δ*Moget3*, Δ*Moget4* and Δ*Mosgt2* strains. After 12, 16 and 24 h post-inoculation (hpi), we peeled the epidermal cells and observed for the development of invasive hyphae. All the strains breached the host cuticle after 12 h of infection, with Δ*Moget3* mutant developing more hyphal branches after 16 and 24 hpi (Fig 6B).

### MoGet1 and MoGet2 are required for hydrophobin synthesis

Since the Δ*Moget1/*Δ*Moget2* mutant strain was unable to form appressoria on both artificial and natural hydrophobic surfaces, we speculated that failure to form appressorium could be attributed to its inability to sense and attach to their host’s hydrophobic surface due to lack of hydrophobin secretion [49–52]. To unravel this puzzle, we dropped 10 µl each of hydrophobicity testing solutions (ddH_2_O, 0.2% gelatin, 0.2% sodium dodecyl sulfate (SDS) + 50 mM EDTA) on the colony surfaces of Δ*Moget1/*Δ*Moget2* and Guy11 stains and incubated for 5 minutes. Results showed a retention of spherical shaped droplets solution on Guy11 mycelia, but collapsed and spread on the Δ*Moget1/*Δ*Moget2* mutant mycelia (Fig 7A). This result suggests a compromise in hydrophobin secretion in the Δ*Moget1/*Δ*Moget2* mutant. Furthermore, we checked the expression of genes involved in hydrophobin synthesis: *MoMPG1* (MGG_10315), *MoMHP1* (MGG_07047), MGG_10105 and MGG_09134 [52–54] in both Guy11 and the Δ*Moget1/*Δ*Moget2* mutant. The expression of the four genes were significantly downregulated in Δ*Moget1*/Δ*Moget*2 mutant compared to their expressions in Wild type Guy11 strain (Fig 7B). Taken together, this result shows that MoGET1 and MoGET2 regulate hydrophobin biosynthesis in *M. oryzae*.

**Fig 7.**
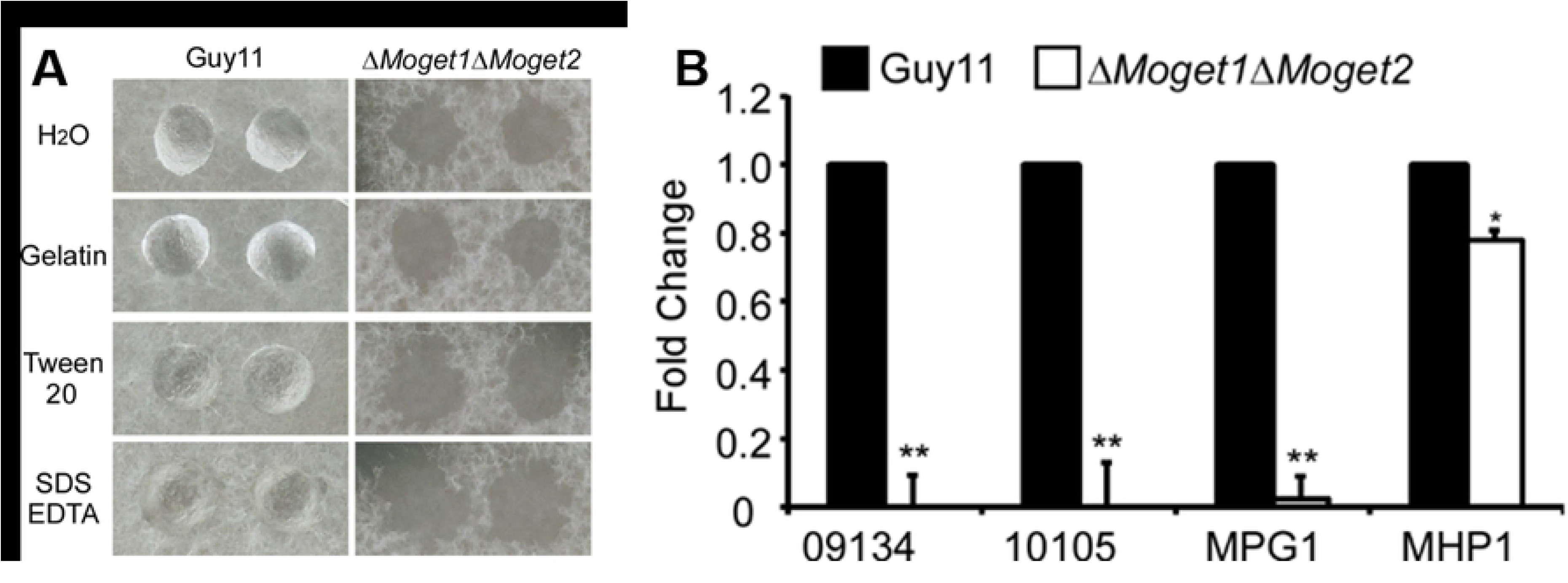
Detergent wettable phenotype of Guy11 and Δ*Moget1/*Δ*Moget2* in hydrophobicity solutions. (A) Hydrophobicity test on the hyphae of Guy11 and Δ*Moget1/*Δ*Moget2* in hydrophobicity test solutions. Surface hydrophobicity of the wild-type strain Guy11 and Δ*Moget1*/Δ*Moget2* mutant was assessed by placing 10 µl each of the test solution (ddH_2_O, 0.2% gelatine, 0.2% sodium dodecyl sulfate (SDS) + 50 mM EDTA, or 250 µg/ml Tween20) on the colony of the strains. The photographs were taken after 5 minutes. The sample solution containing Δ*Moget1/*Δ*Moget*2 double mutant hyphae collapsed while that of Guy11 remained hydrophobic after 5 minutes. (B) Relative expression of hydrophobin-encoding genes, *MoMPG1*, *MoMHP1* and two *MoMHP1* homologues (MGG_09134 and MGG_10105) in the wild type strain Guy11 and Δ*Moget1*/Δ*Moget2* double mutant. Error bars represent SD from the mean of three independent replicates and asterisks represent significant difference between Guy11 and Δ*Moget1*//Δ*Moget2* mutant (** *p* < 0.01).

### MoGet1/MoGet2 complex regulates the autolysis and phosphorylation of Mps1

Natural self-digestion of fungal mycelia normally occurs in aged filamentous fungal cultures. In our analysis of colony morphology of the strains, we observed autolysis in Δ*Moget1/*Δ*Moget2* culture after 6 days of incubation (Fig 8A). This suggests that MoGet1and MoGet2 jointly regulate ageing and and preceding autolysis in *M. oryzae*.

**Fig 8.**
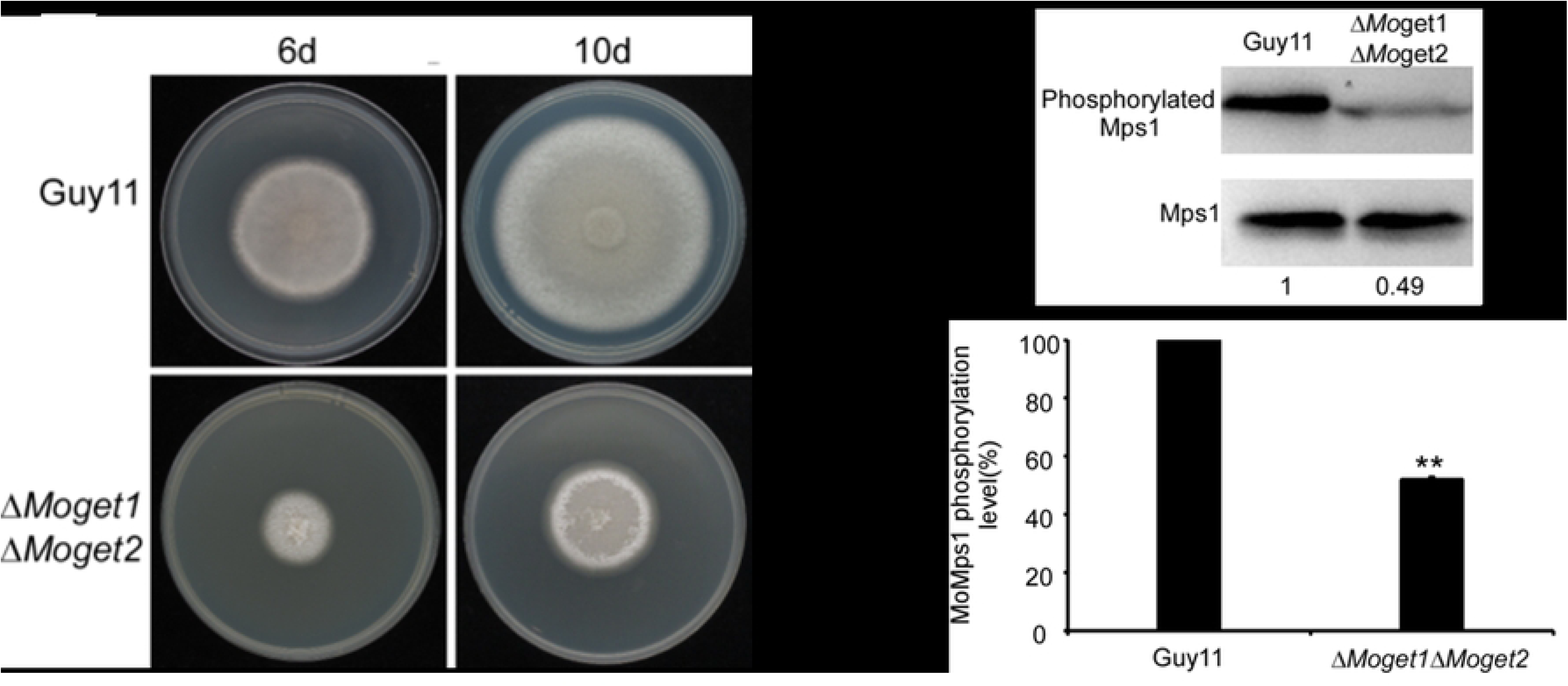
Autolysis and phosphorylation signal of Δ*Moget1/*Δ*Moget2*. (A) Guy11 wild-type and Δ*Moget1/*Δ*Moget2* cultured on CM II for 6 and 10 days. After 10 days, Δ*Moget1/*Δ*Moget1* mycelial mass undergo heavy self-digestion at the distal region, unlike the other strains, suggesting that MoGet1 and MoGet2 act synergistically to regulate autolysis. The complex could be a target for drug discovery to bring about the death of the fungus. (B) Western blot analysis of Mps1 MAPK phosphorylation in Guy11 and Δ*Moget1/*Δ*Moget2* double mutant. Phosphorylation of MoMps1 in Guy11 and Δ*Moget1/Moget2* was detected by p44/42 MAPK (Erk1/2) antibody. Total Proteins were prepared from Guy11 and Δ*Moget1/*Δ*moget2* mutant mycelia cultured in liquid CM medium. The phosphorylation level of MpMps1 in Δ*Moget1/*Δ*moget2* was significantly reduced by about 50% as depicted under the band. (C) Analysis of MoMps1 phosphorylation in Guy11 and the mutant strain. The phosphorylation level was quantified by scanning computer image analysis system using Tanon-5200 (Shanghai Co. Ltd, China).

Furthermore, Mps1 MAPK signaling cascade in *M. oryzae* is a popular pathway known to regulate cell wall biogenesis, fungal development and pathogenesis [55], and the activation of the pathway occurs through phosphorylation of MoMps1 [56]. To further investigate the roles of MoGet1/Moget2 complex in *M. oryzae* development and determine whether the MoGet1 and MoGet2 are directly involved in Mps1 activation, we checked the phosphorylation level of Mps1 in the mutants. A western blot analysis revealed that, compared to the wild type, the phosphorylation level of Mps1 was significantly reduced by about 50% in the *ΔMoget1/*Δ*Moget2* mutant (Fig 8B and 8C), indicating that MoGet1/Moget2 plays an important synergistic role in regulating MoMps1 activation.

### MoGet1 and MoGet2 are localized to the ER while MoGet3, MoGet4 and MoSgt2 are cytosolic

To investigate the resident organelle of each of the five proteins under study, we fused *MoGET1*, *MoGET2*, *MoGET3*, *MoGET4* and *MoSGT2* along with their respective native promoters to a pYF11-GFP plasmid containing geneticin resistant gene at the C-terminal. The constructs were transformed into the protoplasts of the respective mutants and selected on TB3 solid media supplemented with G418 sulfate. The fluorescence microscopy results showed that MoGet1 and MoGet2 were evenly distributed to a particular organelle while MoGet3, MoGet4 and MoSgt2 were clearly localized to the cytoplasm (Fig 9B). To ascertain the actual organelle to which MoGet1 and MoGet2 are localized, we transformed RFP-HDEL ER Marker into the strains and observe their hyphae and conidia under a laser scanning confocal microscope. We found that the GFP and RFP signals colocalized in both the hyphae and conidia (Fig 9A), suggesting the localization of MoGet1 and MoGet2 to the ER of *M. oryzae* and is supported by transmembrane domain that exist in the two proteins (Fig 1A and 1B).

**Fig 9.**
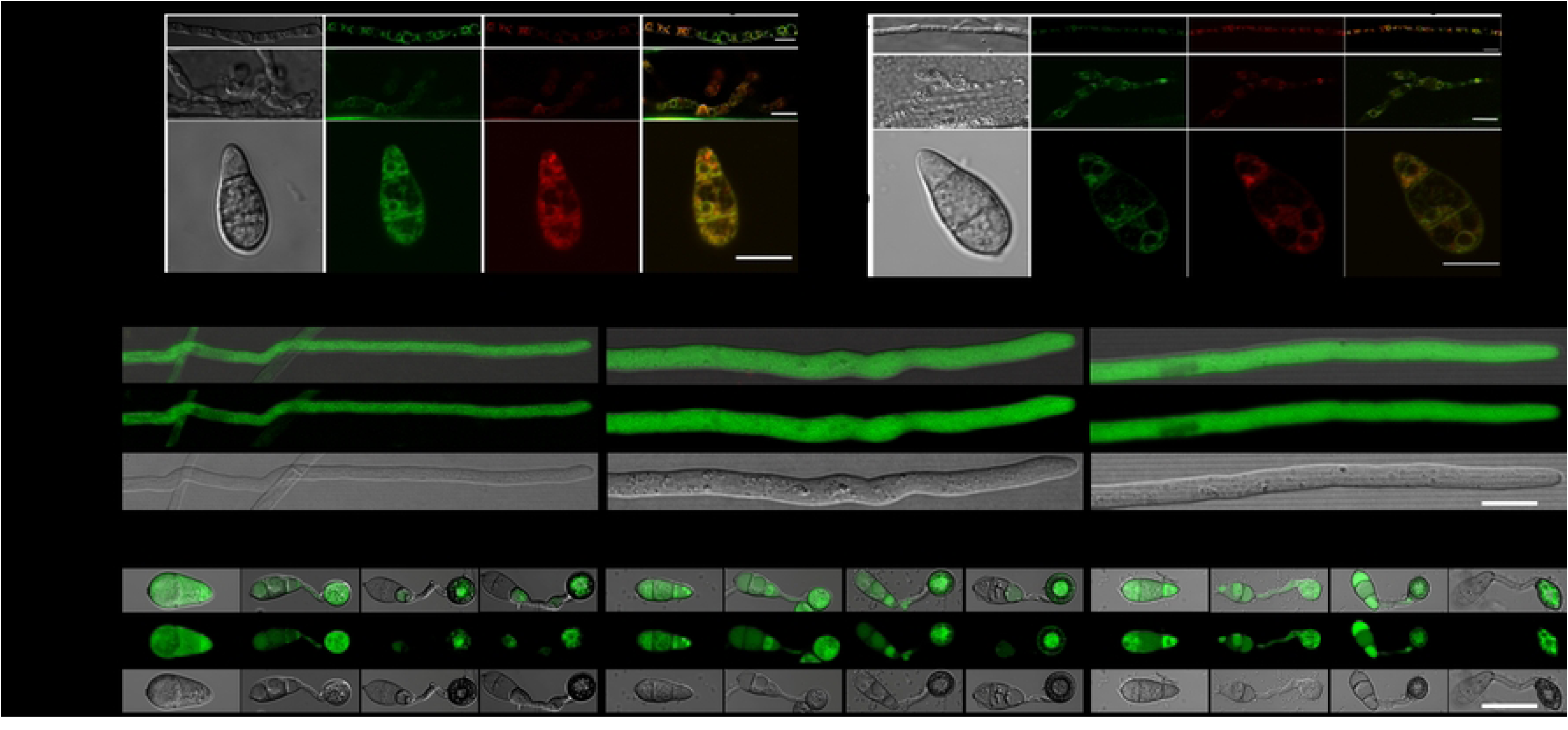
Subcellular localisation of MoGet1, MoGet2, MoGet3, MoGet4 and MoSgt2 at different stages of development. (A) Co-localisation pattern of MoGet1 and MoGet2 in hyphae and conidia of *M. oryzae*. The MoGet1- and MoGet2-GFP do colocalize with the ER marker RFP-HDEL. (B) Localisation of cytosolic MoGet3, MoGet4 and MoSgt2 in the hyphae, conidia and appressorium of *Magnaporthe oryzae*. The live cell images were taken using Nikon Air Laser confocal microscopy. Error bar = 10 µm.

### Y2H and co-immunoprecipitation assays suggest interaction among MoGet1, MoGet2 and MoGet3

The GFP and RFP fluorescence indicated that MoGet1 and MoGet2 were mainly localized to the ER (Fig 1A). Therefore, we hypothesized that the high expression and residence of MoGet1 and MoGet2 in the ER membrane could be due to existence of interaction between the two proteins. To test this hypothesis, we investigated the possible interaction between MoGet1 and MoGet2 or MoGet3 in *M. oryzae* via yeast two hybrid (Y2H) and Co-IP assays. The Y2H analysis showed that MoGet1 interacts with MoGet2 and MoGet3 (Fig 10B). Similarly, MoGet3 also showed positive interaction with MoGet2. Consistently, the results of the Co-IP assay confirmed that MoGet1 and MoGet2 interact with each other and with MoGet3 protein (Fig 10C). Therefore, we conclude that MoGet1 have direct relationship with MoGet2 and MoGet3, and likely forms a complex with MoGet2 on the ER membrane since MoGet1 and MoGet2 have transmembrane domain, unlike MoGet3 that lacks the domain (Fig 1).

**Fig 10.**
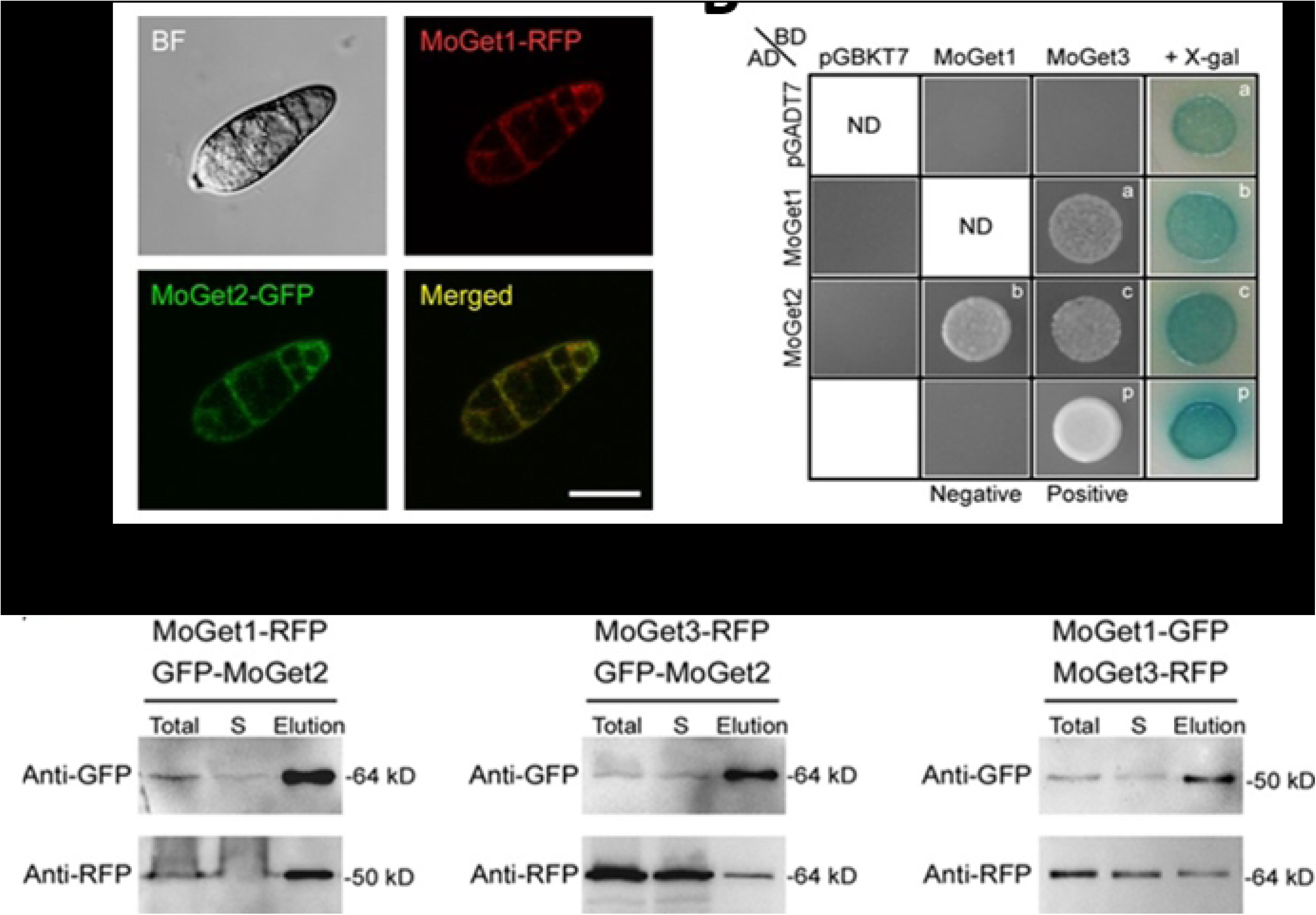
Yeast two hybrid and co-immunoprecipitation assay of MoGet1, MoGet2 and MoGet3. (A) Confocal Laser Scanning showing co-localisation of MoGet1 and MoGet2 GFP-RFP in spore of *M. oryzae*. Merge and individual fluorescence channels are shown. (B) Interaction between MoGet1 and MoGet2 and/or MoGet3 in rice blast fungus. (C) Western blot analysis of Co-Immunoprecipitated MoGet1, MoGet2 and MoGet3 tagged with GFP.

## Discussion

Rice blast disease, caused by the filamentous fungus *Magnaporthe oryzae,* is one of the major diseases of rice (*Oryza sativa*). Functional analysis of genes responsible for blast disease pathogenesis is crucial for the development of mitigation strategy.

In recent years, the structural, biochemical and functional characteristics of GET pathway components (such as Get1, Get2, Get3, Get4, Get5 and Sgt2) have been well studied in model yeast cell (*S. cerevisiae*) [16, 57–59], mammals (*Rattus*) [28, 60], *Plasmodium falciparum* [14] and *Arabidopsis thaliana* [61–63]. However, their functions in filamentous fungal pathogen *M. oryzae* is unknown. This study deployed functional genomic analysis and biochemical approach to elucidate the role of the five Get proteins (Get1, Get2, Get3, Get4, and Sgt2) in *M. oryzae*. Our domain architecture analysis shows that MoGet1 and MoGet2 have transmembrane domain in contrast to MoGet3 and MoGet4, agreeing with the finding in yeast that ScGet3 and ScGet4 are cytosolic and do not require transmembrane domain [64–66]. However, unlike MoGet1, ScGet1 lack transmembrane domain (Fig 1A). MoSgt2 domain architecture, on the other hand, has TPR domain, which is conserved across all the strains analyzed. The TPR domain interacts with Ssa1 in *S. cerevisiae* to recruit TA proteins from the ribosome [67].

Upon deletion, our phenotypic functional analysis study showed that MoGet1 and MoGet2 are indispensable for vegetative growth and virulence of *M. oryzae*. The knockout mutants Δ*Moget1* and Δ*Moget2* were significantly retarded on CM, CM II, and all other solid media used for growth assay in this study, with Δ*Moget2* lacking aerial mycelia, befitting the vital function of TA protein insertion [68]. However, Δ*Moget1* exceeded Guy11 in CM broth, the reason for which will be ascertained through further investigation. The reduced growth exhibited by Δ*Moget1* on solid media is similar to the previous findings in *Arabidopsis thaliana* where root hair elongation was impaired in Get1-deficient *Atget1*mutant strain[69]. This distorted vegetative growth could be attributed to TA mislocalization, toxicity of aggregated TA proteins in the cytosol, deficient mitophagy, or ER stress, as previously demonstrated in yeast cells and *Arabidopsis thaliana* [70, 71]. Although there is no clear report of a direct effect of Get1 and Get2 on filamentation in yeast cells, Zhang *et al* demonstrated that Gets-client protein Scs2 was essential for vegetative growth, asexual development and pathogenicity of *M. oryzae* [72]. Δ*Moget3,* on the other hand, had more expanded vegetative growth than Guy11 after 10 days of incubation. The increased vegetative growth displayed by Δ*Moget3* contrast with the previous finding in *Arabidopsis thaliana,* where the deletion of *AtGET3* resulted in the loss of root hairs and reduced growth

Sporulation is critical in propagating and perpetuating the disease cycle of phytopathogenic fungi. In this study, our asexual reproduction analysis of each deletion mutant strain showed that Moget1, Moget2 and their double deletion mutant are defective in conidiation. The mutant strains failed to grow conidiophores or produce conidia (Fig 2D and Table 1) after repeated attempts using spore-inducing media and conditions. However, the re-introduction of MoGET1 MoGET2 into the genome of the respective mutant strains stores the Guy11-like conidiation. Therefore, we conclude that ER-resident MoGet1 and MoGet2 proteins [73] (Fig 9A) are essential for asexual reproduction of *M. oryzae*. Unexpectedly, the loss of MoGet3 led to an increase in spore production by *M. oryzae*, suggesting that MoGet3 is a repressor of asexual reproduction in the rice blast fungus. This result is contrary to what was obtained in mammals, where the loss of TRC40/ASNA-1, Get3 ortholog in mammals led to embryonic lethality in mice [74] and impaired growth and insulin secretion in *Caenorhabditis elegans* [75].

In eukaryotes, osmotic stress leads to a response necessary for adaptation to hyperosmotic environments [76]. The adaptability of fungus and its propagules to diverse environmental changes (for instance, drought) and conditions is necessary for its survival and proliferation. In this study, our osmotic response test of GET complex factors established that mycelial growth of*Δ*Δ*Moget2* was significantly (p > 0.001) inhibited in media supplemented with osmotic stressors 1 M KCl, 1 M NaCl and 1 M Sorbitol compared to Guy11 wild-type control (Fig 3A and Table 2). However, Δ*Moget1/*Δ*Moget2*, Δ*Moget3*, Δ*Moget4* and Δ*Mosgt2* assumed almost equal inhibition rate as the wild-type, suggesting MoGet2, but MoGet1, MoGet3, MoGet4 and MoSgt2, contributes to osmotic and K^+^ and Na^+^ ionic stress response of *M. oryzae*, and MoGet1 is required for *ΔMoget2*-induced osmotic stress response. On sorbitol-supplemented CM II medium, Δ*Moget1* deletion mutant had its inhibition rate significantly (p > 0.001) reduced compared to Guy11 wild-type (Fig 3B), suggesting that MoGet1 and MoGet2 act antagonistically in the GETs pathway to regulate sorbitol-mediated stress response in *M. oryzae*.

The fungal cell wall contains many of the pathogen-associated molecular patterns (PAMP) (for instance, chitin and β-glucan) recognized by host pattern recognition receptors (PRRs) [77, 78] during host-pathogen interactions. During the interactions, one of the defence responses of the host plant is the release of ROS as an antimicrobial compound to restrict fungal invasion [79, 80]. Although pathogenic fungi have evolved techniques to perceive and neutralize host-secreted enzymatic ROS [79], the overproduction of ROS (oxidative bust) inhibits biotrophic hyphal growth by causing localized cell death around the inoculation site [80]. In this study, our ROS stress response assay result established that Δ*Moget1/Moget2* was significantly inhibited on CM II media supplemented with 5mM H_2_O_2_ ROS compared to its two single mutants, Guy11 and the other strains (Fig 3), suggesting that Δ*Moget1/Moget2* growth defect and failure to cause infection on the host could be, partly, attributed to its inability to sense and detoxify plant-produced ROS in the early stage of infection. It can be inferred that MoGet1 and MoGet2 are interdependent and involved in aiding the adaptation of *M. oryzae* to H_2_O_2_ stressor. The result demonstrates epistatic interactions in which the function of one gene is completely dependent on the presence of a second one [81, 82]. This result is similar and expands on the previous finding in *S. cerevisiae,* where Δ*fus1/*Δ*fus2*, Δ*fus1/*Δ*spa1*, Δ*fus1/*Δ*rvs161* and Δ*fus2/*Δ*spa2* double mutants exhibited stronger cell fusion defects than their four single mutants [83].

Studies on genes involved in pathogenicity is essential for our overall knowledge of disease processes, and any of such genes could be a target for disease control and management [84]. After disrupting the GETs genes, our pathogenicity assay on the host barley and rice leaves using spores or mycelial plugs established that both Δ*Moget1* and Δ*Moget2,* as well as their double knockout mutant Δ*Moget1/*Δ*Moget2,* are non-pathogenic on both wounded and unwounded host tissues (Fig 5A and 5B). The mutants failed to cause the characteristic blast lesions; however, the pathogenicity defects of *M. oryzae* resulting from the disruption of *MoGET1* and *MoGET2* were restored in the complemented strains. More so, our penetration assay indicated that these mutants could develop penetration hyphal into the primary host cell but failed to branch into invasive hyphae (Fig 6A) after 1 – 3 weeks post-inoculation, suggesting that *MoGET1* and *MoGET2* are involved in invasion and pathogenicity. Δ*Moget3*, on the other hand, is more pathogenic than Guy11 wild-type. The penetration assay showed that Δ*Moget3* penetration peg could branch into invasive hyphae within 16 hpi, unlike the wild-type (Fig 6B). Increased virulence exhibited by ΔMoget3 mimics previous finding where Δ*MoaclR* and Δ*Mobzip3* were more pathogenic than the wild type strain [85, 86].

Hydrophobin coating mediates the attachment of fungi to host hydrophobic surfaces during host-pathogen interactions [53, 87]. It can be suggested that Δ*Moget1/*Δ*Moget2* failed in appressorium formation due to inability to produced Hydrophobin (Fig 7B). The participation of the MoGet1 and Moget2 in hydrophobin synthesis/hyphal hydrophobicity [87] may involve direct/indirect interaction between the proteins and the hydrophobic-related pathway. Further investigation is required to ascertain the crosstalk between hydrophobin genes and MoGet1/MoGet2 insertase [71] in *M. oryzae*.

The fungal cell wall is essential for fungal morphology, adaptation, viability, integrity, strength and rigidity against external environmental stress factors, and the internal turgor pressure (0.2 – 10 MPa) that generates the mechanical force required for the penetrability of vegetative hyphal tips through stiff host cuticles [48, 88, 89]. The assembly of its components (mainly chitin, glucan and galactosaminogalactan, GAG) are essential processes that involve a range of fungal-specific enzymes and regulatory networks, therefore making fungal cell walls an attractive target for fungicide as host cells lack many of the cell wall-related proteins [90]. Deleting the fungal cell integrity gene leads to mutants showing defective growth and pathogenicity. Accordingly, our functional analysis result of Get proteins revealed that Δ*Moget2* was significantly (p < 0.0001) inhibited on CFW cell wall stressor compared to the Guy11. Although the growth of Δ*Moget2* on CR-supplemented media is more retarded than Guy11, the inhibition rate is less significant compared to what we obtained on media supplemented with CFW (Fig 4A and Table 3). More so, Δ*Moget1* and Δ*Moget2* are highly sensitive to ER-stress-inducing reducing agent dithiothreitol (DTT) compared to Guy11 wild-type and the other strains, suggesting that MoGet1 and MoGet2 are required for *M. oryzae* ER membrane integrity. Intriguingly, the Δ*Moget1/*Δ*Moget2* double deletion mutant had its inhibition rate significantly (p > 0.001) reduced compared to its single two mutant strains (Table 3). We hypothesize that the disruption of MoGET1/MoGET2 resulted in a new ER-membrane integrity protein(s) replacing the MoGet1/MoGet2 complex. Further research is needed to investigate and unravel this mystery. By contrast, DTT had only minor to no effect on Δ*Moget3*, Δ*Moget4* and Δ*Mosgt2* conforming with their cytosolic nature. The least inhibition of Δ*Moget3* by CR and CFW compared to Guy11 (Table 3) shows that MoGet3 is a negative regulator of cell wall integrity. More so, the near equal inhibition rate of Guy11, the mutants and their complemented strains on media supplemented with cell membrane stressor, SDS, indicates that the GETs component factors are not essential for cell membrane integrity.

Furthermore, there were more protoplasts released from Δ*Moget2* mutant than Guy11 (Fig 4B and 4C), after treatment with lysing enzyme at two different time points: 30 min and 1 h 30 min, confirming further the poor and compromised cell wall integrity of MoGet2-deficient strain.

The MoGet1 and MoGet2 completely colocalized to the ER, while MoGet3, MoGet4, and MoSgt2 are localized to the cytoplasm. The co-localization of MoGet1 and MoGet2 to the ER is evident in the sensitivity of MoGet1- and MoGet2-deficient mutants to ER stressor DDT and supports a synergistic role in the recruitment of Get3-TA protein complex to the ER lipid bilayer [43]. This result agrees with the previous finding, which established that *S. cerevisiae* ScGet1 and ScGet2 are ER-resident [71, 91] while Moget3, MoGet4 and MoSgt2 are cytosolic [27, 92]. But, our domain architecture analysis showed that, unlike MoGet1, MoGet2 and ScGet2, ScGet1 lacks a transmembrane domain (Fig 1A and 1B). This observation suggests that while MoGet1 and MoGet2 reside and jointly recruit TA proteins into the ER lipid bilayer of *M. oryzae*, only ScGet2 of *S. cerevisiae* reside in ER, with ScGet1 possibly hanging in the cytosol to recruit TA protein substrates from ScGet3 and transfer it to ScGet2 in the *S. cerevisiae* ER membrane.

Most biological processes, including transcription regulation and signal transduction within and between cells, are governed by interactions between proteins [93], and the phenotype of an organism may be the combinational output of multiple input signals. In this study, our Y2H assay and Co-IP analysis of MoGet1-GFP, GFP-MoGet2 and MoGet3-RFP (Fig 10A) demonstrated that MoGet1, MoGet2 and MoGet3 strongly interact with one another (Fig 10B and 10C), similar to earlier findings in *S. cerevisiae* and *Arabidopsis thaliana* [16, 43, 69]. In *S. cerevisiae*, this interaction and the delivery of TA protein is ATP-dependent [16]. However, how these interactions occur in *M. oryzae* is yet unknown, so further investigation is needed.

In summary, we have demonstrated that GET complex components are conserved in *M. oryzae* and perform unique functions in vegetative growth, hydrophobicity, conidiation and pathogenicity. While MoGet4 and MoSgt2 have no significant function in *M. oryzae* pathogenicity, MoGet1 and MoGet2 are required for vegetative growth, conidiation and pathogenicity. We also demonstrated that MoGet2 is essential for cell wall integrity, osmotic and ER stress response and that MoGet1 and MoGet2 work synergistically to respond to H_2_O_2_ ROS stress. MoGet3, on the other hand, is a negative regulator of vegetative growth, cell wall integrity, conidiation and pathogenicity in *M. oryzae*. Moreover, we established that MoGet1 and MoGet2 colocalized to the ER and interacted with each other as well as with MoGet3. However, the live cell imaging of fluorescently labeled Get proteins indicates that MoGet3, MoGet4 and MoSgt2 are localized to the cytoplasm.

Since MoGet3, MoGet4 and MoSgt2 are dispensable for vegetative growth, sporulation, appressorium formation, invasive growth and pathogenicity of *M. oryzae,* we suggest there is an alternative route for TA proteins from, possibly Ssa1, to ER membrane. This may be so because TA proteins must be transferred to their destination organelle to carry out their essential physiological functions of vesicular trafficking, protein translocation across organelle, programmed cell death, protein quality control and organelle dynamics and tethering [94]. Therefore, further research is needed to unravel the probable alternative pathway for TA proteins in *M. oryzae*.

## Acknowledgements

We are grateful to Yang Zifeng, Wilfred M. Anjago, Huxiao Xu, Ibrahim Tijjani and Jiyu Su for the helpful project discussions.

## Author contributions

**Conceptualization:** Zonghua Wang and Wei Tang.

**Data curation:** Wei Tang and Felix Abah.

**Formal analysis:** Felix Abah and Qiaojia Zheng.

**Funding acquisition:** Zonghua Wang.

**Investigation:** Felix Abah and Qiaojia Zheng.

**Methodology:** Wei Tang, Felix Abah, Qiaojia Zheng and Linwan Huang, Xiaomi Chen and Jules Biregeya.

**Project administration:** Zonghua Wang, Wei Tang and Felix Abah.

**Resources:** Zonghua Wang.

**Supervision:** Zonghua Wang and Wei Tang.

**Validation:** Zonghua Wang, Wei Tang and Felix Abah.

**Wrting – original draft:** Felix Abah.

**Writing – review and editing:** Aron Osakina and Felix Abah.

## Supporting information

**S1 Fig. Confirmation of single insertion of hygromycin Phosphotransferase gene (*HPH*) in ORF region by Southern blot analysis.** **A.** Probe1, Probe2, Probe3, Probe4 and Probe5 correspond to *MoGET1*, *MoGET2*, *MoGET3*, *MoGET4* and *MoSGT2* gene coding regions probe, respectively. **B.** Southern blot analysis of *M. oryzae* Guy11 wild- type and the mutant strains. Genomic DNA extracted from wild-type control and the mutants were digested with *Hind*III and hybridized with probe specific for *HPH* fragment.

**S1 Table. Primers used in this study.**

## References

1. Asibi AE, Chai Q, Coulter JA. Rice Blast: A Disease with Implications for Global Food Security. Agronomy. 2019;9(8):451. PubMed PMID: doi:10.3390/agronomy9080451.

2. Muthayya S, Sugimoto JD, Montgomery S, Maberly GF. An overview of global rice production, supply, trade, and consumption. Annals of the New York Academy of Sciences. 2014;1324:7–14. Epub 2014/09/17. doi: 10.1111/nyas.12540. PubMed PMID: 25224455.

3. Samal P, Babu SC, Mondal B, Mishra SN. The global rice agriculture towards 2050: An inter-continental perspective. Outlook on Agriculture. 2022:00307270221088338.

4. Osés-Ruiz M, Cruz-Mireles N, Martin-Urdiroz M, Soanes DM, Eseola AB, Tang B, et al. Appressorium-mediated plant infection by *Magnaporthe oryzae* is regulated by a Pmk1-dependent hierarchical transcriptional network. Nature microbiology. 2021;6(11):1383–97.

5. Nalley L, Tsiboe F, Durand-Morat A, Shew A, Thoma G. Economic and Environmental Impact of Rice Blast Pathogen (*Magnaporthe oryzae*) Alleviation in the United States. PloS one. 2016;11(12):e0167295. Epub 2016/12/03. doi: 10.1371/journal.pone.0167295. PubMed PMID: 27907101; PubMed Central PMCID: PMCPMC5131998.

6. Longya A, Talumphai S, Jantasuriyarat C. Morphological Characterization and Genetic Diversity of Rice Blast Fungus, *Pyricularia oryzae*, from Thailand Using ISSR and SRAP Markers. Journal of fungi (Basel, Switzerland). 2020;6(1). Epub 2020/03/25. doi: 10.3390/jof6010038. PubMed PMID: 32204416; PubMed Central PMCID: PMCPMC7151035.

7. Cruz-Mireles N, Eseola AB, Osés-Ruiz M, Ryder LS, Talbot NJ. From appressorium to transpressorium—Defining the morphogenetic basis of host cell invasion by the rice blast fungus. PLoS pathogens. 2021;17(7):e1009779.

8. Wilson RA. Magnaporthe oryzae. Trends in microbiology. 2021;29(7):663–4.

9. Oliveira-Garcia E, Tamang TM, Park J, Dalby M, Martin-Urdiroz M, Rodriguez Herrero C, et al. Clathrin-mediated endocytosis facilitates the internalization of *Magnaporthe oryzae* effectors into rice cells. The Plant cell. 2023;35(7):2527–51. Epub 2023/03/29. doi: 10.1093/plcell/koad094. PubMed PMID: 36976907; PubMed Central PMCID: PMCPMC10291035.

10. Rocha RO, Elowsky C, Pham NTT, Wilson RA. Spermine-mediated tight sealing of the *Magnaporthe oryzae* appressorial pore-rice leaf surface interface. Nature microbiology. 2020;5(12):1472–80. Epub 2020/09/16. doi: 10.1038/s41564-020-0786-x. PubMed PMID: 32929190.

11. Wang M, Dean RA. Host induced gene silencing of *Magnaporthe oryzae* by targeting pathogenicity and development genes to control rice blast disease. Frontiers in plant science. 2022;13:959641. Epub 2022/08/30. doi: 10.3389/fpls.2022.959641. PubMed PMID: 36035704; PubMed Central PMCID: PMCPMC9403838.

12. Aron O, Wang M, Lin L, Batool W, Lin B, Shabbir A, et al. MoGLN2 Is Important for Vegetative Growth, Conidiogenesis, Maintenance of Cell Wall Integrity and Pathogenesis of *Magnaporthe oryzae*. Journal of fungi (Basel, Switzerland). 2021;7(6). Epub 2021/07/03. doi: 10.3390/jof7060463. PubMed PMID: 34201222; PubMed Central PMCID: PMCPMC8229676.

13. Rada P, Makki A, Žárský V, Tachezy J. Targeting of tail-anchored proteins to *Trichomonas vaginalis* hydrogenosomes. Molecular microbiology. 2019;111(3):588–603. Epub 2018/12/07. doi: 10.1111/mmi.14175. PubMed PMID: 30506591.

14. Kumar T, Maitra S, Rahman A, Bhattacharjee S. A conserved guided entry of tail-anchored pathway is involved in the trafficking of a subset of membrane proteins in *Plasmodium falciparum*. PLoS pathogens. 2021;17(11):e1009595. Epub 2021/11/16. doi: 10.1371/journal.ppat.1009595. PubMed PMID: 34780541; PubMed Central PMCID: PMCPMC8629386.

15. Borgese N, Brambillasca S, Colombo S. How tails guide tail-anchored proteins to their destinations. Current opinion in cell biology. 2007;19(4):368–75. Epub 2007/07/17. doi: 10.1016/j.ceb.2007.04.019. PubMed PMID: 17629691.

16. Schuldiner M, Metz J, Schmid V, Denic V, Rakwalska M, Schmitt HD, et al. The GET complex mediates insertion of tail-anchored proteins into the ER membrane. Cell. 2008;134(4):634–45. Epub 2008/08/30. doi: 10.1016/j.cell.2008.06.025. PubMed PMID: 18724936; PubMed Central PMCID: PMCPMC2572727.

17. Pedrazzini E. Tail-anchored proteins in plants. Journal of Plant Biology. 2009;52:88–101.

18. Borgese N, Colombo S, Pedrazzini E. The tale of tail-anchored proteins: coming from the cytosol and looking for a membrane. The Journal of cell biology. 2003;161(6):1013–9. Epub 2003/06/25. doi: 10.1083/jcb.200303069. PubMed PMID: 12821639; PubMed Central PMCID: PMCPMC2173004.

19. Hegde RS, Keenan RJ. Tail-anchored membrane protein insertion into the endoplasmic reticulum. Nature reviews Molecular cell biology. 2011;12(12):787–98. Epub 2011/11/17. doi: 10.1038/nrm3226. PubMed PMID: 22086371; PubMed Central PMCID: PMCPMC3760496.

20. Jonikas MC, Collins SR, Denic V, Oh E, Quan EM, Schmid V, et al. Comprehensive characterization of genes required for protein folding in the endoplasmic reticulum. Science (New York, NY). 2009;323(5922):1693-7. Epub 2009/03/28. doi: 10.1126/science.1167983. PubMed PMID: 19325107; PubMed Central PMCID: PMCPMC2877488.

21. Shan SO. Guiding tail-anchored membrane proteins to the endoplasmic reticulum in a chaperone cascade. The Journal of biological chemistry. 2019;294(45):16577–86. Epub 2019/10/03. doi: 10.1074/jbc.REV119.006197. PubMed PMID: 31575659; PubMed Central PMCID: PMCPMC6851334.

22. Hegde RS, Keenan RJ. The mechanisms of integral membrane protein biogenesis. Nature reviews Molecular cell biology. 2022;23(2):107–24. Epub 2021/09/25. doi: 10.1038/s41580-021-00413-2. PubMed PMID: 34556847.

23. Asseck LY, Mehlhorn DG, Monroy JR, Ricardi MM, Breuninger H, Wallmeroth N, et al. Endoplasmic reticulum membrane receptors of the GET pathway are conserved throughout eukaryotes. Proceedings of the National Academy of Sciences of the United States of America. 2021;118(1). Epub 2021/01/15. doi: 10.1073/pnas.2017636118. PubMed PMID: 33443185; PubMed Central PMCID: PMCPMC7817167.

24. Voth W, Schick M, Gates S, Li S, Vilardi F, Gostimskaya I, et al. The protein targeting factor Get3 functions as ATP-independent chaperone under oxidative stress conditions. Molecular Cell. 2014;56(1):116–27.

25. Kohl C, Tessarz P, von der Malsburg K, Zahn R, Bukau B, Mogk A. Cooperative and independent activities of Sgt2 and Get5 in the targeting of tail- anchored proteins. Biological chemistry. 2011;392(7):601–8. Epub 2011/05/31. doi: 10.1515/bc.2011.066. PubMed PMID: 21619481.

26. Chartron JW, Gonzalez GM, Clemons WM, Jr. A structural model of the Sgt2 protein and its interactions with chaperones and the Get4/Get5 complex. The Journal of biological chemistry. 2011;286(39):34325–34. Epub 2011/08/13. doi: 10.1074/jbc.M111.277798. PubMed PMID: 21832041; PubMed Central PMCID: PMCPMC3190793.

27. Chang YW, Chuang YC, Ho YC, Cheng MY, Sun YJ, Hsiao CD, et al. Crystal structure of Get4-Get5 complex and its interactions with Sgt2, Get3, and Ydj1. The Journal of biological chemistry. 2010;285(13):9962–70. Epub 2010/01/29. doi: 10.1074/jbc.M109.087098. PubMed PMID: 20106980; PubMed Central PMCID: PMCPMC2843242.

28. Colombo SF, Cardani S, Maroli A, Vitiello A, Soffientini P, Crespi A, et al. Tail-anchored Protein Insertion in Mammals: FUNCTION AND RECIPROCAL INTERACTIONS OF THE TWO SUBUNITS OF THE TRC40 RECEPTOR. The Journal of biological chemistry. 2016;291(29):15292–306. Epub 2016/05/27. doi: 10.1074/jbc.M115.707752. PubMed PMID: 27226539; PubMed Central PMCID: PMCPMC4946941.

29. Liu W, Xie Y, Ma J, Luo X, Nie P, Zuo Z, et al. IBS: an illustrator for the presentation and visualization of biological sequences. 2015;31(20):3359–61.

30. 30. Kumar S, Stecher G, Tamura K. MEGA7: Molecular Evolutionary Genetics Analysis Version 7.0 for Bigger Datasets. Mol Biol Evol. 2016;33(7):1870–4. Epub 2016/03/24. doi: 10.1093/molbev/msw054. PubMed PMID: 27004904; PubMed Central PMCID: PMCPMC8210823.

31. Abah F, Kuang Y, Biregeya J, Abubakar YS, Ye Z, Wang Z. Mitogen-Activated Protein Kinases SvPmk1 and SvMps1 Are Critical for Abiotic Stress Resistance, Development and Pathogenesis of *Sclerotiophoma versabilis*. Journal of fungi (Basel, Switzerland). 2023;9(4). Epub 2023/04/28. doi: 10.3390/jof9040455. PubMed PMID: 37108909; PubMed Central PMCID: PMCPMC10142639.

32. Lin L, Cao J, Du A, An Q, Chen X, Yuan S, et al. eIF3k Domain-Containing Protein Regulates Conidiogenesis, Appressorium Turgor, Virulence, Stress Tolerance, and Physiological and Pathogenic Development of *Magnaporthe oryzae* Oryzae. Frontiers in plant science. 2021:2316.

33. He S, Huang K, Li B, Lu G, Wang A. Functional Analysis of a Salicylate Hydroxylase in *Sclerotinia sclerotiorum*. Journal of fungi (Basel, Switzerland). 2023;9(12). Epub 2023/12/22. doi: 10.3390/jof9121169. PubMed PMID: 38132770; PubMed Central PMCID: PMCPMC10744347.

34. Kuang Y-b, Abah F, Abubakar YS, Ye Z-y, Wang Z-h. Establishment of protoplast preparation protocol and genetic transformation system for *Sclerotiophoma versabilis*. Journal of Plant Pathology. 2023:1–9.

35. Cai Y, Liu X, Shen L, Wang N, He Y, Zhang H, et al. Homeostasis of cell wall integrity pathway phosphorylation is required for the growth and pathogenicity of *Magnaporthe oryzae*. Molecular plant pathology. 2022;23(8):1214–25. Epub 2022/05/05. doi: 10.1111/mpp.13225. PubMed PMID: 35506374; PubMed Central PMCID: PMCPMC9276948.

36. Lin L, Cao J, Du A, An Q, Chen X, Yuan S, et al. eIF3k Domain-Containing Protein Regulates Conidiogenesis, Appressorium Turgor, Virulence, Stress Tolerance, and Physiological and Pathogenic Development of *Magnaporthe oryzae* Oryzae. Frontiers in plant science. 2021;12:748120. Epub 2021/11/05. doi: 10.3389/fpls.2021.748120. PubMed PMID: 34733303; PubMed Central PMCID: PMCPMC8558559.

37. Anjago WM, Biregeya J, Shi M, Chen Y, Wang Y, Wang Z, et al. The Calcium Chloride Responsive Type 2C Protein Phosphatases Play Synergistic Roles in Regulating MAPK Pathways in *Magnaporthe oryzae*. Journal of fungi (Basel, Switzerland). 2022;8(12). Epub 2022/12/23. doi: 10.3390/jof8121287. PubMed PMID: 36547620; PubMed Central PMCID: PMCPMC9784850.

38. Zheng H, Zheng W, Wu C, Yang J, Xi Y, Xie Q, et al. Rab GTP ases are essential for membrane trafficking - dependent growth and pathogenicity in Fusarium graminearum. 2015;17(11):4580–99.

39. Anjago WM, Biregeya J, Shi M, Chen Y, Wang Y, Wang Z, et al. The calcium chloride responsive type 2C protein phosphatases play synergistic roles in regulating MAPK pathways in Magnaporthe oryzae. 2022;8(12):1287.

40. Bruno KS, Tenjo F, Li L, Hamer JE, Xu JR. Cellular localization and role of kinase activity of PMK1 in *Magnaporthe grisea*. Eukaryotic cell. 2004;3(6):1525–32. Epub 2004/12/14. doi: 10.1128/ec.3.6.1525-1532.2004. PubMed PMID: 15590826; PubMed Central PMCID: PMCPMC539019.

41. Dean R, Van Kan JA, Pretorius ZA, Hammond-Kosack KE, Di Pietro A, Spanu PD, et al. The Top 10 fungal pathogens in molecular plant pathology. Molecular plant pathology. 2012;13(4):414–30. Epub 2012/04/05. doi: 10.1111/j.1364-3703.2011.00783.x. PubMed PMID: 22471698; PubMed Central PMCID: PMCPMC6638784.

42. Carvalho HJF, Del Bondio A, Maltecca F, Colombo SF, Borgese N. The WRB Subunit of the Get3 Receptor is Required for the Correct Integration of its Partner CAML into the ER. Scientific reports. 2019;9(1):11887. Epub 2019/08/17. doi: 10.1038/s41598-019-48363-2. PubMed PMID: 31417168; PubMed Central PMCID: PMCPMC6695381.

43. Wang F, Chan C, Weir NR, Denic V. The Get1/2 transmembrane complex is an endoplasmic-reticulum membrane protein insertase. Nature. 2014;512(7515):441-4. Epub 2014/07/22. doi: 10.1038/nature13471. PubMed PMID: 25043001; PubMed Central PMCID: PMCPMC4342754.

44. Batool W, Shabbir A, Lin L, Chen X, An Q, He X, et al. Translation Initiation Factor eIF4E Positively Modulates Conidiogenesis, Appressorium Formation, Host Invasion and Stress Homeostasis in the Filamentous Fungi *Magnaporthe oryzae*. Frontiers in plant science. 2021;12:646343. Epub 2021/07/06. doi: 10.3389/fpls.2021.646343. PubMed PMID: 34220879; PubMed Central PMCID: PMCPMC8244596.

45. Duran R, Cary JW, Calvo AM. Role of the osmotic stress regulatory pathway in morphogenesis and secondary metabolism in filamentous fungi. Toxins. 2010;2(4):367–81. Epub 2010/04/01. doi: 10.3390/toxins2040367. PubMed PMID: 22069590; PubMed Central PMCID: PMCPMC3153207.

46. Blomberg A, Adler L. Physiology of osmotolerance in fungi. Advances in microbial physiology. 1992;33:145–212.

47. Ray PD, Huang BW, Tsuji Y. Reactive oxygen species (ROS) homeostasis and redox regulation in cellular signaling. Cellular signalling. 2012;24(5):981–90. Epub 2012/01/31. doi: 10.1016/j.cellsig.2012.01.008. PubMed PMID: 22286106; PubMed Central PMCID: PMCPMC3454471.

48. Garcia-Rubio R, de Oliveira HC, Rivera J, Trevijano-Contador N. The Fungal Cell Wall: *Candida*, *Cryptococcus*, and *Aspergillus Species*. Frontiers in microbiology. 2019;10:2993. Epub 2020/01/30. doi: 10.3389/fmicb.2019.02993. PubMed PMID: 31993032; PubMed Central PMCID: PMCPMC6962315.

49. Weber G, Bornscheuer UT, Wei R. Enzymatic Plastic Degradation: Academic Press; 2021.

50. Bernard K. The Genus Corynebacterium Reference Module in Biomedical Sciences 2016. doi: 10.1016/b978-0-12-801238-3.99200-6

51. Dubey MK, Jensen DF, Karlsson M. Hydrophobins are required for conidial hydrophobicity and plant root colonization in the fungal biocontrol agent *Clonostachys rosea*. BMC microbiology. 2014;14:18. Epub 2014/02/04. doi: 10.1186/1471-2180-14-18. PubMed PMID: 24483277; PubMed Central PMCID: PMCPMC3922079.

52. Kim S, Ahn IP, Rho HS, Lee YH. MHP1, a *Magnaporthe grisea* hydrophobin gene, is required for fungal development and plant colonization. Molecular microbiology. 2005;57(5):1224–37.

53. Talbot NJ, Kershaw MJ, Wakley GE, De Vries O, Wessels J, Hamer JE. MPG1 Encodes a Fungal Hydrophobin Involved in Surface Interactions during Infection-Related Development of *Magnaporthe grisea*. The Plant cell. 1996;8(6):985–99. Epub 1996/06/01. doi: 10.1105/tpc.8.6.985. PubMed PMID: 12239409; PubMed Central PMCID: PMCPMC161153.

54. Yan Y, Wang H, Zhu S, Wang J, Liu X, Lin F, et al. The methylcitrate cycle is required for development and virulence in the rice blast fungus Pyricularia oryzae. 2019;32(9):1148–61.

55. Zhou T, Dagdas YF, Zhu X, Zheng S, Chen L, Cartwright Z, et al. The glycogen synthase kinase MoGsk1, regulated by Mps1 MAP kinase, is required for fungal development and pathogenicity in Magnaporthe oryzae. 2017;7(1):945.

56. Jiang C, Zhang X, Liu H, Xu J-RJPp. Mitogen-activated protein kinase signaling in plant pathogenic fungi. 2018;14(3):e1006875.

57. Josefson R, Kumar N, Hao X, Liu B, Nyström T. The GET pathway is a major bottleneck for maintaining proteostasis in *Saccharomyces cerevisiae*. Scientific reports. 2023;13(1):9285. Epub 2023/06/08. doi: 10.1038/s41598-023-35666-8. PubMed PMID: 37286562; PubMed Central PMCID: PMCPMC10247811.

58. Onishi M, Kubota M, Duan L, Tian Y, Okamoto K. The GET pathway serves to activate Atg32-mediated mitophagy by ER targeting of the Ppg1-Far complex. Life science alliance. 2023;6(4). Epub 2023/01/26. doi: 10.26508/lsa.202201640. PubMed PMID: 36697253; PubMed Central PMCID: PMCPMC9880027.

59. Denic V, Dötsch V, Sinning I. Endoplasmic reticulum targeting and insertion of tail-anchored membrane proteins by the GET pathway. Cold Spring Harbor perspectives in biology. 2013;5(8):a013334. Epub 2013/08/03. doi: 10.1101/cshperspect.a013334. PubMed PMID: 23906715; PubMed Central PMCID: PMCPMC3721280.

60. Farkas Á, Bohnsack KE. Capture and delivery of tail-anchored proteins to the endoplasmic reticulum. The Journal of cell biology. 2021;220(8). Epub 2021/07/16. doi: 10.1083/jcb.202105004. PubMed PMID: 34264263; PubMed Central PMCID: PMCPMC8287540.

61. Srivastava R, Zalisko BE, Keenan RJ, Howell SHJPp. The GET system inserts the tail-anchored protein, SYP72, into endoplasmic reticulum membranes. 2017;173(2):1137–45.

62. Anderson SA, Satyanarayan MB, Wessendorf RL, Lu Y, Fernandez DEJTPC. A homolog of GuidedEntry of Tail-anchored proteins3 functions in membrane- specific protein targeting in chloroplasts of Arabidopsis. 2021;33(8):2812–33.

63. Mehlhorn DG, Asseck LY, Grefen CJPP. Looking for a safe haven: tail- anchored proteins and their membrane insertion pathways. 2021;187(4):1916–28.

64. Powis K, Schrul B, Tienson H, Gostimskaya I, Breker M, High S, et al. Get3 is a holdase chaperone and moves to deposition sites for aggregated proteins when membrane targeting is blocked. Journal of cell science. 2013;126(Pt 2):473–83. Epub 2012/12/04. doi: 10.1242/jcs.112151. PubMed PMID: 23203805; PubMed Central PMCID: PMCPMC3613179.

65. Chang YW, Lin TW, Li YC, Huang YS, Sun YJ, Hsiao CD. Interaction surface and topology of Get3-Get4-Get5 protein complex, involved in targeting tail- anchored proteins to endoplasmic reticulum. The Journal of biological chemistry. 2012;287(7):4783–9. Epub 2011/12/23. doi: 10.1074/jbc.M111.318329. PubMed PMID: 22190685; PubMed Central PMCID: PMCPMC3281643.

66. Simpson PJ, Schwappach B, Dohlman HG, Isaacson RL. Structures of Get3, Get4, and Get5 provide new models for TA membrane protein targeting. Structure (London, England : 1993). 2010;18(8):897–902. Epub 2010/08/11. doi: 10.1016/j.str.2010.07.003. PubMed PMID: 20696390; PubMed Central PMCID: PMCPMC3557799.

67. Shan SO. Role of Hsp70 in Post-Translational Protein Targeting: Tail-Anchored Membrane Proteins and Beyond. Int J Mol Sci. 2023;24(2). Epub 2023/01/22. doi: 10.3390/ijms24021170. PubMed PMID: 36674686; PubMed Central PMCID: PMCPMC9866221.

68. Abell BM, Mullen RTJPcr. Tail-anchored membrane proteins: exploring the complex diversity of tail-anchored-protein targeting in plant cells. 2011;30:137–51.

69. Xing S, Mehlhorn DG, Wallmeroth N, Asseck LY, Kar R, Voss A, et al. Loss of GET pathway orthologs in *Arabidopsis thaliana* causes root hair growth defects and affects SNARE abundance. Proceedings of the National Academy of Sciences of the United States of America. 2017;114(8):E1544–e53. Epub 2017/01/18. doi: 10.1073/pnas.1619525114. PubMed PMID: 28096354; PubMed Central PMCID: PMCPMC5338382.

70. Srivastava R, Zalisko BE, Keenan RJ, Howell SH. The GET System Inserts the Tail-Anchored Protein, SYP72, into Endoplasmic Reticulum Membranes. Plant physiology. 2017;173(2):1137-45. Epub 2016/12/08. doi: 10.1104/pp.16.00928. PubMed PMID: 27923985; PubMed Central PMCID: PMCPMC5291014.

71. Onishi M, Nagumo S, Iwashita S, Okamoto K. The ER membrane insertase Get1/2 is required for efficient mitophagy in yeast. Biochemical and biophysical research communications. 2018;503(1):14–20. Epub 2018/04/21. doi: 10.1016/j.bbrc.2018.04.114. PubMed PMID: 29673596.

72. Zhang J, Chen X, Yang Z, Xu H, Weng S, Wang Z, et al. Endoplasmic reticulum membrane protein MoScs2 is important for asexual development and pathogenesis of *Magnaporthe oryzae*. Frontiers in microbiology. 2022;13:906784. Epub 2022/08/23. doi: 10.3389/fmicb.2022.906784. PubMed PMID: 35992683; PubMed Central PMCID: PMCPMC9386004.

73. Rivera-Monroy J, Musiol L, Unthan-Fechner K, Farkas Á, Clancy A, Coy- Vergara J, et al. Mice lacking WRB reveal differential biogenesis requirements of tail-anchored proteins in vivo. Scientific reports. 2016;6:39464. Epub 2016/12/22. doi: 10.1038/srep39464. PubMed PMID: 28000760; PubMed Central PMCID: PMCPMC5175141.

74. Mukhopadhyay R, Ho YS, Swiatek PJ, Rosen BP, Bhattacharjee H. Targeted disruption of the mouse Asna1 gene results in embryonic lethality. FEBS letters. 2006;580(16):3889–94. Epub 2006/06/27. doi: 10.1016/j.febslet.2006.06.017. PubMed PMID: 16797549.

75. Kao G, Nordenson C, Still M, Rönnlund A, Tuck S, Naredi P. ASNA-1 positively regulates insulin secretion in *C. elegans* and mammalian cells. Cell. 2007;128(3):577–87. Epub 2007/02/10. doi: 10.1016/j.cell.2006.12.031. PubMed PMID: 17289575.

76. Hohmann S. Control of high osmolarity signalling in the yeast *Saccharomyces cerevisiae*. FEBS letters. 2009;583(24):4025–9. Epub 2009/11/03. doi: 10.1016/j.febslet.2009.10.069. PubMed PMID: 19878680.

77. Vega K, Kalkum M. Chitin, chitinase responses, and invasive fungal infections. International journal of microbiology. 2012;2012:920459. Epub 2011/12/22. doi: 10.1155/2012/920459. PubMed PMID: 22187561; PubMed Central PMCID: PMCPMC3236456.

78. Zhang H, Yang J, Liu M, Xu X, Yang L, Liu X, et al. Early molecular events in the interaction between Magnaporthe oryzae and rice. 2024;6(1):9.

79. Zhang N, Lv F, Qiu F, Han D, Xu Y, Liang W. Pathogenic fungi neutralize plant-derived ROS via Srpk1 deacetylation. The EMBO journal. 2023;42(9):e112634. Epub 2023/03/10. doi: 10.15252/embj.2022112634. PubMed PMID: 36891678; PubMed Central PMCID: PMCPMC10152141.

80. Singh Y, Nair AM, Verma PK. Surviving the odds: From perception to survival of fungal phytopathogens under host-generated oxidative burst. Plant communications. 2021;2(3):100142. Epub 2021/05/25. doi: 10.1016/j.xplc.2021.100142. PubMed PMID: 34027389; PubMed Central PMCID: PMCPMC8132124.

81. 81. Miko IJNE. Epistasis: gene interaction and phenotype effects. 2008;1(1):197.

82. van Leeuwen J, Boone C, Andrews BJ. Mapping a diversity of genetic interactions in yeast. Current opinion in systems biology. 2017;6:14–21. Epub 2018/12/07. doi: 10.1016/j.coisb.2017.08.002. PubMed PMID: 30505984; PubMed Central PMCID: PMCPMC6269142.

83. Gammie AE, Brizzio V, Rose MDJMbotc. Distinct morphological phenotypes of cell fusion mutants. 1998;9(6):1395–410.

84. Idnurm A, Howlett BJ. Pathogenicity genes of phytopathogenic fungi. Molecular plant pathology. 2001;2(4):241–55. Epub 2001/07/01. doi: 10.1046/j.1464-6722.2001.00070.x. PubMed PMID: 20573012.

85. Liu C, Shen N, Zhang Q, Qin M, Cao T, Zhu S, et al. *Magnaporthe oryzae* Transcription Factor MoBZIP3 Regulates Appressorium Turgor Pressure Formation during Pathogenesis. Int J Mol Sci. 2022;23(2). Epub 2022/01/22. doi: 10.3390/ijms23020881. PubMed PMID: 35055065; PubMed Central PMCID: PMCPMC8778449.

86. Osés-Ruiz M, Cruz-Mireles N, Martin-Urdiroz M, Soanes DM, Eseola AB, Tang B, et al. Appressorium-mediated plant infection by *Magnaporthe oryzae* is regulated by a Pmk1-dependent hierarchical transcriptional network. Nature microbiology. 2021;6(11):1383–97. Epub 2021/10/29. doi: 10.1038/s41564-021-00978-w. PubMed PMID: 34707224.

87. Degani O, Lev S, Ronen MJP, pathology mp. Hydrophobin gene expression in the maize pathogen Cochliobolus heterostrophus. 2013;83:25–34.

88. Plaza V, Silva-Moreno E, Castillo L. Breakpoint: Cell Wall and Glycoproteins and their Crucial Role in the Phytopathogenic Fungi Infection. Current protein & peptide science. 2020;21(3):227–44. Epub 2019/09/07. doi: 10.2174/1389203720666190906165111. PubMed PMID: 31490745.

89. Gow NAR, Latge JP, Munro CA. The Fungal Cell Wall: Structure, Biosynthesis, and Function. Microbiology spectrum. 2017;5(3). Epub 2017/05/18. doi: 10.1128/microbiolspec.FUNK-0035-2016. PubMed PMID: 28513415.

90. Zhao C-R, You Z-L, Chen D-D, Hang J, Wang Z-B, Ji M, et al. Structure of a fungal 1, 3-β-glucan synthase. 2023;9(37):eadh7820.

91. Zhang Y, Schäffer T, Wölfle T, Fitzke E, Thiel G, Rospert S. Cotranslational Intersection between the SRP and GET Targeting Pathways to the Endoplasmic Reticulum of *Saccharomyces cerevisiae*. Molecular and cellular biology. 2016;36(18):2374–83. Epub 2016/06/30. doi: 10.1128/mcb.00131-16. PubMed PMID: 27354063; PubMed Central PMCID: PMCPMC5007794.

92. Wang F, Whynot A, Tung M, Denic V. The mechanism of tail-anchored protein insertion into the ER membrane. Molecular cell. 2011;43(5):738–50. Epub 2011/08/13. doi: 10.1016/j.molcel.2011.07.020. PubMed PMID: 21835666; PubMed Central PMCID: PMCPMC3614002.

93. Lu H-C, Fornili A, Fraternali FJErop. Protein–protein interaction networks studies and importance of 3D structure knowledge. 2013;10(6):511–20.

94. Jiang H. Quality control pathways of tail-anchored proteins. Biochimica et biophysica acta Molecular cell research. 2021;1868(2):118922. Epub 2020/12/08. doi: 10.1016/j.bbamcr.2020.118922. PubMed PMID: 33285177.

